# A broad-spectrum phage-encoded mechanism to disarm bacterial type IV filaments

**DOI:** 10.64898/2026.03.10.710608

**Authors:** Nathan Roberge, Paankhi Dave, Véronique Taylor, Taylor J. Ellison, Callum Myers, Courtney K. Ellison, Karen L. Maxwell, Lori L. Burrows

## Abstract

Phages can modify host cell physiology to thwart competitors. The *Pseudomonas aeruginosa*-specific phage DMS3 encodes Aqs1, a protein inhibitor of type IV pilus (T4P) function to prevent host cell recognition by other phages that leverage these filaments for infection. Aqs1 disrupts T4P by binding to the hexameric ATPase PilB, required to power pilus filament extension, though several mechanistic details remain unclear. We show that Aqs1 has broad-spectrum activity and can disrupt T4P function in a variety of Gram negative bacteria. This protein inhibits PilB by binding to a solvent-exposed hydrophobic patch on the N2-domain, distal to the active site. Binding destabilizes the hexamer, preventing PilB accumulation at T4P machines. Aqs1 likely disrupts PilB oligomerization by displacing a flexible linker segment between the PilB N1- and N2-domains required for inter-subunit contact. Together, the Aqs1 mode of action provides a design template for broad-spectrum inhibitors of diverse bacterial virulence factors.

**Significance:** The phage-encoded protein Aqs1 disables type IV pilus (T4P) production in *Pseudomonas aeruginosa* by targeting the hexameric ATPase responsible for assembling pilus fibers. We show that despite originating from a *P. aeruginosa*-specific phage, Aqs1 can also disable T4 ATPase-dependent phenotypes across other pathogenic bacteria and homologous systems. Mechanistically, Aqs1 binds to a conserved patch on the PilB N2-domain away from the active site. Binding here breaks apart the PilB oligomer, preventing it from acting on T4P machines. Aqs1 binding at the N2-domain patch likely displaces a flexible PilB linker segment that binds to this site to stabilize the hexamer. Our work highlights a novel and conserved PilB allosteric site which is exploited by the phage-encoded protein Aqs1 to disrupt diverse T4 systems in multiple bacteria.

## Introduction

Bacterial viruses, known as phages, hijack host cellular machinery to replicate and disseminate throughout a population^1^. During infection, some phages will also modify host cell surface architecture and/or physiology^2–5^. For example, proteins expressed by certain phages can disable the production of type IV pili (T4P) in the pathogenic bacterium *Pseudomonas aeruginosa*^2,3,6^. This helps to evade recognition by related phages which use these structures as receptors to infect cells^2,7^. T4P are important virulence factors, enabling cells to interact with surfaces, sense contact, and migrate across them to form biofilms^8–12^. In other bacteria, they also facilitate uptake of DNA from the environment which can contribute to antimicrobial resistance^13,14^. Related complexes like the type 2 secretion system (T2SS) instead are used for extracellular export of protein toxins during host colonization^15^. Some phage-encoded proteins inhibit pilus function by targeting the homohexameric ATPase, PilB, in the cytoplasm at the base of the T4P nanomachine^16–18^. PilB, in conjunction with additional protein regulatory effectors, is an essential component of T4P function, responsible for driving assembly and extension of pilus filaments into extracellular space^19–22^. How many of these phage-encoded proteins interact with and inhibit PilB is poorly understood, but since all T4P systems rely on the activity of PilB-like ATPases^23^, understanding this mechanism could provide a useful template for the design of anti-virulence therapeutics.

One of these phage-encoded proteins, anti-quorum sensing protein 1 (Aqs1), expressed by the pilus-targeting phage DMS3, was recently shown to inhibit both PilB and the principal quorum sensing regulator LasR through distinct interfaces^2^. A separate study found that an Aqs1 homologue called twitching inhibitory protein (Tip) disrupted accumulation of PilB at the cell poles^3^, where T4P are produced in this organism^18,24,25^, but neither investigation explained these findings mechanistically. Here, we show that Aqs1 provides broad inhibition of T4P-dependent natural competence and surface motility by inhibiting PilB in *Acinetobacter baylyi* and *Stenotrophomonas maltophilia*, respectively. It also inhibits the *P. aeruginosa* T2SS likely by targeting GspE, the conserved PilB homologue for this system. Aqs1 binds to a hydrophobic allosteric patch on the PilB N2-domain, distal to the ATP-binding active site^26–28^. Binding there disrupts PilB oligomerization and polar localization, likely by displacing a flexible linker segment located between the N-terminal N1- and N2-domains.

## Results

### Aqs1 can bind to PilB homologues from many species and disrupt their functions

The phage-encoded protein Aqs1 is expressed from DMS3 following entry into *P. aeruginosa*, where it binds to the transcriptional regulator LasR and to PilB (**Fig 1A**). Aqs1-dependent inhibition of each target is mediated through separate surfaces of the protein^2^. Introducing Y39S and R40G substitutions (Aqs1^YR:SG^) to the Aqs1 <u>YR</u>DALD motif critical for interaction with LasR^2^ is sufficient to allow pyocyanin production, consistent with the empty vector control strain (**Fig 1B**) as reported for certain PAO1 isolates^29,30^. The Aqs1^YR:SG^ mutant still blocks T4P-dependent twitching motility and prevents infection by the pilus-targeting phage DMS3 (**Fig 1B**, **C**). Likewise, expression of Aqs1 F19A or W45A mutants that cannot interact with PilB^2^ prevents pyocyanin production but does not impact twitching motility or confer phage resistance (**Fig 1B**, **C**). Aqs1 and Aqs1^YR:SG^ interact with PilB when co-expressed in an *Escherichia coli* bacterial two-hybrid assay (BACTH), but only wild-type Aqs1 interacts with LasR^31^ (**Fig S1**). Notably, expression of neither Aqs1 nor Aqs1^YR:SG^ alters *P. aeruginosa* growth (**Fig S1**) suggesting that the loss of each phenotype stems from selective target inhibition rather than through non-specific action on central pathways.

**Figure 1:**
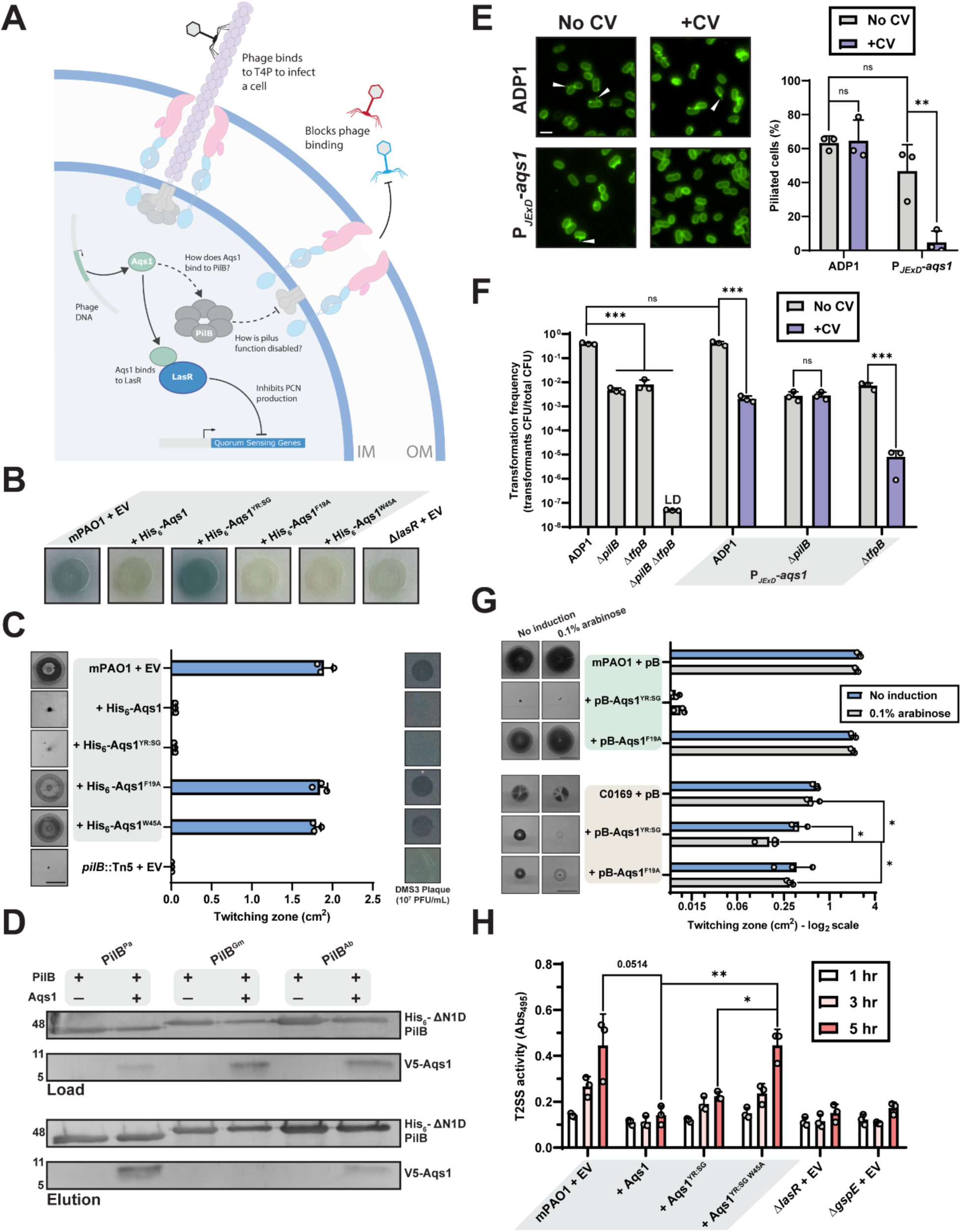
Aqs1 inhibits T4 ATPase-dependent phenotypes from multiple systems and bacteria. **(A)** Phage-encoded protein Aqs1 is expressed following DMS3 entry into *P. aeruginosa*. Aqs1 inhibits LasR-dependent production of pyocyanin and T4P function likely through interaction with PilB. **(B)** Representative colony pyocyanin production (blue) of mPAO1 expressing various His_6_-Aqs1 mutants. **(C)** Quantification of sub-agar stab twitching motility zones of mPAO1 expressing various His_6_-Aqs1 mutants. Representative crystal violet-stained twitching zones are shown to the left. Bars represent the means of triplicate samples from three independent experiments ± SD. Scale bar = 1 cm. Boxes to the right show representative DMS3 plaquing on a bacterial lawn of the same strain. Strains grouped in grey boxes indicate the same background **(D)** Western blot of sample input (Load) and following co-purification (Elution) of N1-domain truncated (ΔN1D)-His_6_-PilB homologues from *P. aeruginosa* (PilB^Pa^), *G. metallireducens* (PilB^Gm^), and *A. baylyi* (PilB^Ab^) with V5-Aqs1. Molecular weight is indicated to the left. Conditions grouped in the grey boxes indicate the same ΔN1D-His_6_-PilB construct. **EV**: empty pHERD30T vector, **His_6_-Aqs1**: N-terminally His_6_-tagged Aqs1 in pHERD30T vector, **His_6_-Aqs1^YR:SG^:** N-terminally His_6_-tagged Aqs1 Y39S R40G in pHERD30T vector, **His_6_-Aqs1^F19A^:** N-terminally His_6_-tagged Aqs1^F19A^ in pHERD30T vector, **His_6_-Aqs1^W45A^:** N-terminally tagged His_6_-tagged Aqs1^W45A^ in pHERD30T vector. All expression was induced with 0.1% arabinose. **(E)** Representative images of indicated strains with labeled T4P grown with or without crystal violet inducer (CV). White arrows indicate visible pili. Scale bar = 2 µm. Number of cells producing visible pili in the indicated populations expressed as a percentage of total cells is shown to the right. Bars represent the means ± SD of three independent experiments. For each biological replicate, a minimum of 50 total cells were assessed. **(F)** Natural transformation assays of indicated strains and growth conditions. Strains in the grey box indicate the same background. Bars represent the means ± SD of three independent experiments. The transformation frequency of the Δ*pilB* Δ*tfpB* strain was below the limit of detection, indicated by LD. **P*_JExD_*-*aqs1*:** His_6_-*aqs1* expressed from the P*_JExD_* inducible promoter. Statistics were determined by Sidak’s multiple comparisons test; log-transformed data were used for statistical analysis for transformation assays. **ns**: not significant; **\*\***: *p* < 0.01; **\*\*\***: *p* < 0.001 **(G)** Quantification of sub-agar stab twitching motility zones of mPAO1 and WCC C0169 expressing Aqs1 mutants. Representative twitching zones are shown to the left. The mPAO1 or C0169 strains grouped in the green and tan boxes respectively indicate the same backgrounds. Bars represent the means of duplicate samples from three independent experiments ± SD. Scale bar = 1 cm. Expression was induced with 0.1% arabinose. **(H)** Quantification of Congo red-elastin cleavage from bacterial culture supernatants at the indicated timepoints. Strains in the grey box indicate the same background. Bars represent the means ± SD of three independent experiments. A full-length multiple sequence alignment of T4P (PilB) and T2SS (GspE) ATPase homologues from *P. aeruginosa* with PilB from *Stenotrophomonas maltophilia* WCC C0169 is shown above. Essential residues for Aqs1 contact are coloured in pink. **\***: 0.05 ≥ *p* ≥ 0.01; **\*\***: 0.01 ≥ *p* ≥ 0.001 (Two-tailed Welch’s *t-*test). **EV**: empty pHERD30T vector, **Aqs1 or indicated mutant:** N-terminally His_6_-tagged Aqs1 or indicated Aqs1 mutant in pHERD30T vector, **pB:** empty pBADGr vector, **pB-Aqs1 or indicated mutant:** Aqs1 or indicated Aqs1 mutant in pBADGr vector.

Many bacteria deploy T4P or homologous systems to engage with and sense surface contact but also to power motility, protein secretion, or take up environmental DNA^8,13–15,32^. The PilB homologues that power these machines are often conserved^28^, yet a PSI-BLAST search for the Aqs core sequence revealed that homologues are not widely distributed beyond the *Pseudomonas* genus. We therefore wondered whether Aqs1 could interact with and disable other similar T4P ATPases outside of this bacterium. To assess this, we co-purified hexahistidine-tagged N-terminal PilB truncations (ΔN1D-PilB) with V5-tagged Aqs1. The N1-domain was truncated to improve PilB stability and solubility as previously described^19,22,33,34^. Under these conditions, PilB homologues from *P. aeruginosa* (PilB^Pa^) and the more conserved *A. baylyi* (PilB^Ab^, 56% pairwise identity; **Fig S2**) eluted with Aqs1 while the less conserved protein from *Geobacter metallireducens* (PilB^Gm^, 43% pairwise identity; **Fig S2**) did not (**Fig 1D**). These data confirmed that Aqs1 can also recognize PilB^Ab^ and thus may impact its function. To test this, we introduced hexahistidine-tagged Aqs1 under the control of a crystal violet (CV)-inducible promoter (P*_JExD_*-*aqs1*) onto the *A. baylyi* ADP1 chromosome and assessed piliation using fluorescence microscopy. Discernable pili were lost upon Aqs1 induction, and we observed a significant reduction in the percentage of piliated cells (**Fig 1E**), consistent with a defect in synthesizing pilus filaments^13,35^. *A. baylyi* encodes two distinct extension ATPases, PilB and TfpB, which can act independently on pilus machines^13^. To test motor specificity, we assessed T4P-dependent DNA uptake in *pilB* and *tfpB* mutants expressing Aqs1. Natural transformation was reduced in the wild type to levels consistent with inhibition of one of the two ATPases (**Fig 1F**). We also observed a significant reduction in transformation frequency in the Δ*tfpB* but not the Δ*pilB* background (**Fig 1F**) indicating that Aqs1 is specific for PilB. Aqs1 does not interact with TfpB in a BACTH assay, further supporting this finding (**Fig S2**). To assess broad disruption of twitching, we expressed Aqs1^YR:SG^ in the cystic fibrosis pathogen *Stenotrophomonas maltophilia* as this species, like *P. aeruginosa*, can deploy pili for motility^36^. While expression in this bacterium conferred a modest growth defect (**Fig S1**), Aqs1^YR:SG^ significantly decreased twitching motility while the non-PilB binding Aqs1^F19A^ variant did not (**Fig 1G**). The *P. aeruginosa* T2SS employs a distinct homologous ATPase, GspE (**Fig S3**), to drive export of toxic protein effectors. We assessed GspE inhibition by quantifying cleavage of a Congo red-elastin conjugated substrate by the secreted protein effector, elastase, in culture supernatants. Secretion is regulated by quorum-sensing, thus expressing Aqs1 abolishes T2SS activity, consistent with deletion of *lasR*^37^ (**Fig 1H**). Substrate cleavage was also significantly reduced, however, in the presence of Aqs1^YR:SG^ but not a non-LasR and PilB binding Aqs1^YR:SG^ ^W45A^ mutant, indicating that T2SS function is impaired independently of LasR inhibition (**Fig 1H**). Taken together, Aqs1 has broad-spectrum activity against PilB-like ATPases in many bacteria.

### Aqs1 inhibits T4P assembly by binding to the N2-domain of PilB

The fact that Aqs1 could disable multiple ATPases across organisms warranted further mechanistic investigation. To understand how Aqs1 interacts with and broadly inhibits PilB function, we modelled the complex using Alphafold3^38^ (**Fig 2A**, **S3**). Aqs1 forms a dimer in solution^2^, but when modelled with the *P. aeruginosa* PilB monomer only a single protomer of the dimer engages the subunit (**Fig 2A**). The bound Aqs1 protomer was predicted to contact the PilB N2-domain distal to the ATP-binding active site, which is formed at the interfaces between the C-terminal domains of each monomer^28^. The side chains of essential Aqs1 residues F19 and W45 were packed against the N2-domain at this putative interface. On PilB, several residues with hydrophobic side chains clustered around the Aqs1 interface, collectively forming a pronounced hydrophobic patch on an otherwise solvent-exposed region of the protein^26–28^ (**Fig 2B**). Several of these residues are conserved in the Aqs1-binding PilB homologues (**Fig 2C**) and ATPases which do not share these sequence features do not interact with Aqs1 (**Fig S2**), collectively supporting this structural prediction. To validate these PilB side chains as targets for Aqs1 binding, we individually mutated each residue to alanine and assessed interaction with Aqs1 in a BACTH assay (**Fig 2D**). Several of the mutants including P228A, P229A, and L232A demonstrated decreased Aqs1 interaction compared to wild-type PilB (**Fig 2D**). Interestingly, only one of the PilB F187A and I236A mutant fusions interacted with Aqs1, which could indicate a lower affinity or transient interaction^31^. Each PilB point mutant complemented twitching motility when expressed in a *pilB* deficient background (**Fig 2E**) confirming that each variant produces a functional gene product.

**Figure 2:**
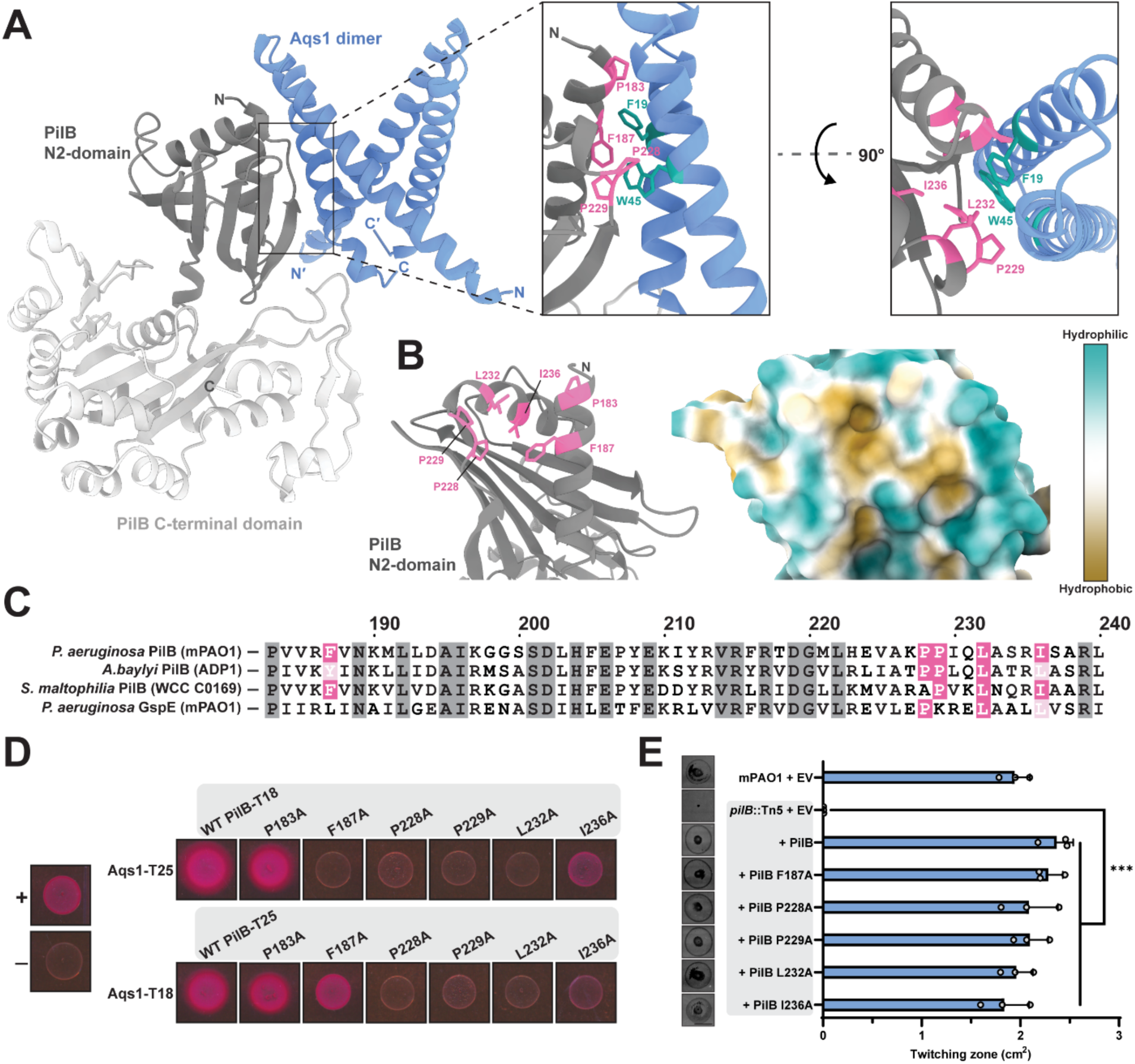
Aqs1 interacts with the PilB N2-domain. **(A)** Alphafold3 model of a full length *P. aeruginosa* PilB monomer with the N-terminal domain and linker region (M1-D180) hidden for clarity, bound to the Aqs1 dimer. The PilB N2-domain and CTD are indicated in dark and light grey respectively. Insets show the putative PilB-Aqs1 interaction interface with residue side chains of interest on PilB shown in pink and Aqs1 F19 and W45 shown in green. **(B)** Alternate view of the PilB N2-domain model with Aqs1 hidden for clarity. Putative Aqs1-interacting side chains are highlighted in pink. Surface hydrophobicity and hydrophilicity overlayed onto the cartoon representation are shown. The indicated PilB side chains form a hydrophobic patch on the N2-domain surface. **(C)** Full length sequence alignment of Aqs1-binding T4 ATPases highlighting the N2-domain. Putative Aqs1-binding residues are shaded in pink. **(D)** Representative colonies showing pairwise interaction (pink) between PilB N2-domain point mutants and Aqs1. Untagged T18/T25 plasmid and PilS-T18/PilS-T25 homodimers were used as negative and positive controls respectively. **(E)** Quantification of sub-agar stab twitching motility zones of mPAO1 *pilB*::Tn5 complemented with each PilB N2-domain mutant. Representative crystal violet-stained twitching zones are shown to the left. Strains grouped in the grey box indicate the same background. Bars represent the means of duplicate samples from three independent experiments ± SD. Scale bar = 1 cm. Expression was induced using 0.05% arabinose. **EV**: empty pHERD30T vector, **PilB or indicated PilB point mutant**: PilB or indicated point mutant in pHERD30T vector. **\*\*\***: 0.001 ≥ *p* (One-way ANOVA).

We individually introduced the F187A, P228A, and P229A mutations onto the *P. aeruginosa* chromosome and assessed T4P-dependent phenotypes when Aqs1^YR:SG^ was expressed. The Aqs1^YR:SG^ mutant was used in subsequent experiments to distinguish between T4P and LasR-dependent phenotypes in *P. aeruginosa*. Expression of Aqs1^YR:SG^ did not inhibit twitching motility of the P228A and P229A mutants (**Fig 3A**) nor block DMS3 infection (**Fig 3B**), indicating that these mutants allowed pili to assemble and were resistant to the activity of Aqs1^YR:SG^. In contrast, expressing Aqs1^YR:SG^ in the F187A background significantly attenuated twitching compared to the empty vector control and only provided partial protection to DMS3 (**Fig 3A, B**), supporting a lower affinity interaction. We next co-expressed wild type hexahistidine-tagged ΔN1D-PilB or the point mutants together with V5-tagged Aqs1 in *E. coli* and performed pull-downs. Under these conditions, wild type PilB co-eluted with Aqs1 from a Ni^2+^-NTA column. The F187A, P228A, and P229A variants pulled down less Aqs1 than the wild type (**Fig 3C**) confirming that, as expected from their phenotypes, these mutants have a lower affinity for Aqs1.

**Figure 3:**
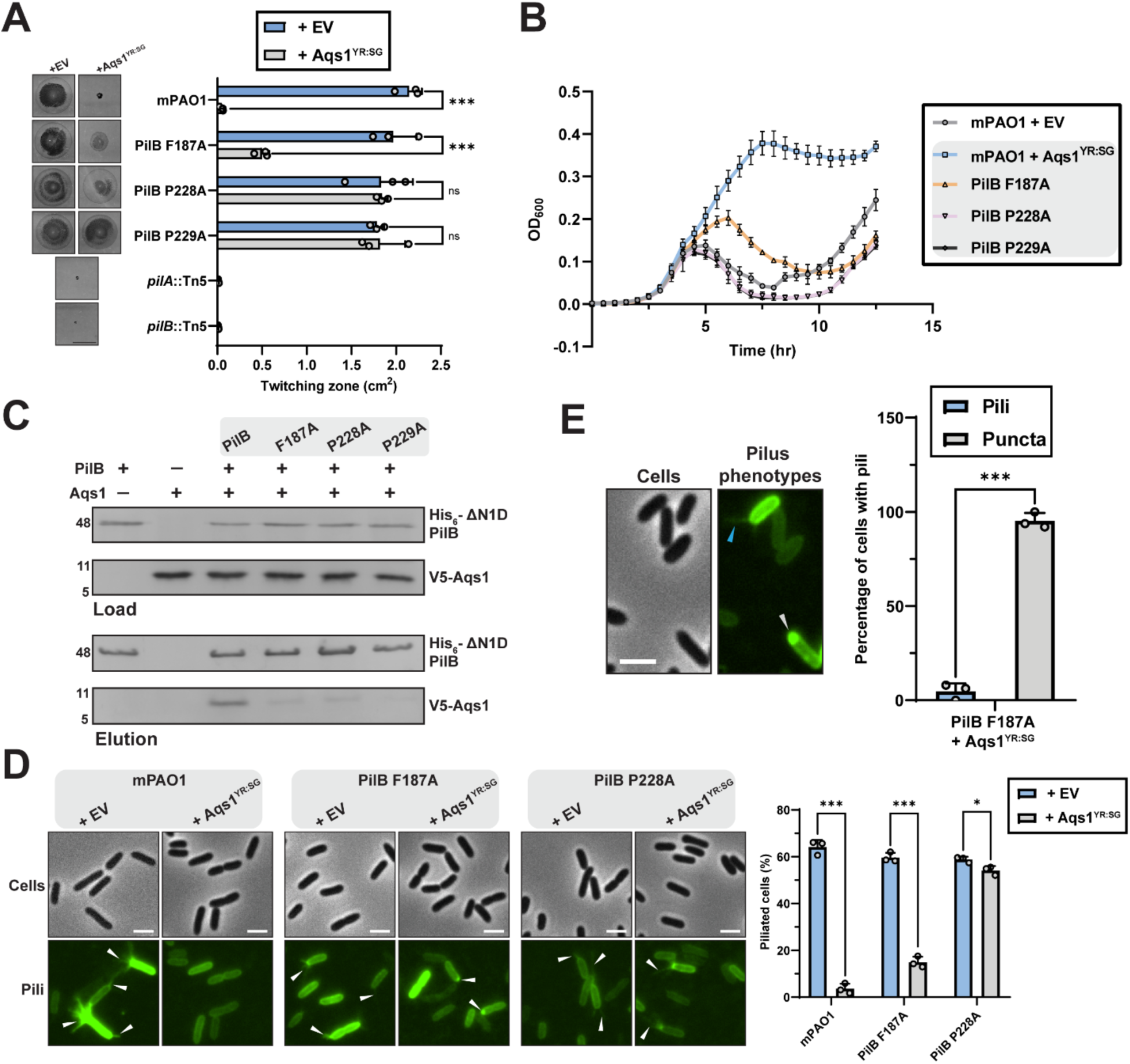
Mutations in the PilB N2-domain reduce Aqs1 interaction and restore T4P phenotypes. **(A)** Quantification of sub-agar stab twitching motility zones of mPAO1 PilB N2-domain chromosomal point mutants expressing His_6_-Aqs1^YR:SG^. Representative crystal violet-stained twitching zones are shown to the left. Bars represent the means of duplicate samples from three independent experiments ± SD. Scale bar = 1 cm. **(B)** Bacterial growth curves across 18 h in LB media of mPAO1 N2-domain point mutants expressing His_6_-Aqs1^YR:SG^ with DMS3 phage-challenge. Points represent the means of triplicate samples from three independent experiments ± SD. **(C)** Western blot of sample input (Load) and following co-purification (Elution) of ΔN1-domain truncated (N1D)-His_6_-PilB variants with V5-Aqs1. Molecular weight is indicated to the left. Indicated point mutations grouped in the grey box are all made in ΔN1D-His_6_-PilB. **(D)** Representative images of PilA A86C type IV pili labelled with maleimide-Alexafluor 488 with background fluorescence subtracted. White arrows indicate visible pili. All scale bars = 2 µm. Strains grouped in the grey boxes indicate the same background. **(E)** Number of cells producing visible PilA A86C type IV pili expressed as a percentage of total cells. Bars represent the means ± SD of three independent experiments. For each biological replicate, a minimum of 50 total cells were assessed. **(F)** Number of piliated PilB F187A cells expressing His_6_-Aqs1^YR:SG^ expressed as a percentage of total cells with discernible polar puncta or visible PilA A86C type IV pili. Bars represent the means ± SD of three independent experiments. **EV**: empty pHERD30T vector, **Aqs1^YR:SG^**: N-terminally His_6_-tagged Aqs1 Y39S R40G in pHERD30T vector. **ns**: *p* ≥ 0.05; **\***: 0.05 ≥ *p* ≥ 0.01; **\*\*\***: 0.001 ≥ *p* (Two-tailed Welch’s *t-*test).

We next assessed the impact of Aqs1^YR:SG^ expression on surface-exposed pilus production by fluorescence microscopy^22,35,39^. The wild type, PilB F187A, and P228A mutants (P229A was not tested since its phenotypes were similar to those of P228A) were similarly piliated (**Fig 3D**), with approximately 60% of the total cells having visible pili (**Fig 3D**). When Aqs1^YR:SG^ was expressed in the wild-type background, only ∼3.5% of cells retained discernable pili (**Fig 3D**). In contrast, the PilB P228A mutant maintained high levels of surface piliation, with only a ∼4% reduction in the percentage of piliated cells in the presence of Aqs1^YR:SG^ (**Fig 3D**). This is consistent with this mutant’s retention of T4P-dependent phenotypes when Aqs1^YR:SG^ is expressed (**Fig 3A**, **B**). In the PilB F187A mutant, approximately 15-20% of cells had bright polarly-associated puncta when Aqs1^YR:SG^ was expressed (**Fig 3D**, **E**), which were scored as an accumulation of short pili^22^. Visible filaments were produced in this background, but these were rare throughout the piliated subpopulation (**Fig 3E**). This could suggest that the lower affinity of Aqs1^YR:SG^ for this mutant dampens PilB inhibition, allowing pilus extension to commence before it is interrupted, consistent with the partial disruption of T4P-phenotypes we observed (**Fig 3A**, **B**). Together, these data support that Aqs1 interacts with PilB at the N2-domain to inhibit its function.

### Aqs1 prevents PilB polar localization by disrupting oligomerization

The Aqs1 homologue Tip was previously reported to delocalize PilB from cell poles through an unknown process^3^. To assess whether Aqs1 acts similarly, we examined PilB subcellular localization in the presence of Aqs1. Tagging PilB can impact its function^24,25^ so we proxied PilB localization using an mNeonGreen-PilZ (mNGr-PilZ) fluorescent fusion expressed from the *P. aeruginosa* chromosome (**Fig 4A**). PilZ is a small chaperone essential for PilB stability and function, and colocalizes with PilB in cells^6,19,20,22^. The PilZ fusion strain was partially functional and Aqs protein expression in this background reduced twitching to *pilB* deficient levels (**Fig S4**). Approximately 30% of control cells showed discernable polar puncta with most cells displaying a unipolar distribution (**Fig 4B**). Expression of Aqs1, Aqs1^YR:SG^, or Tip reduced the percentage of cells with discernable polar puncta to ∼10% in each instance. This was the expected outcome for Tip^3^, but this confirms Aqs1 similarly disrupts localization of pilus extension machinery.

**Figure 4:**
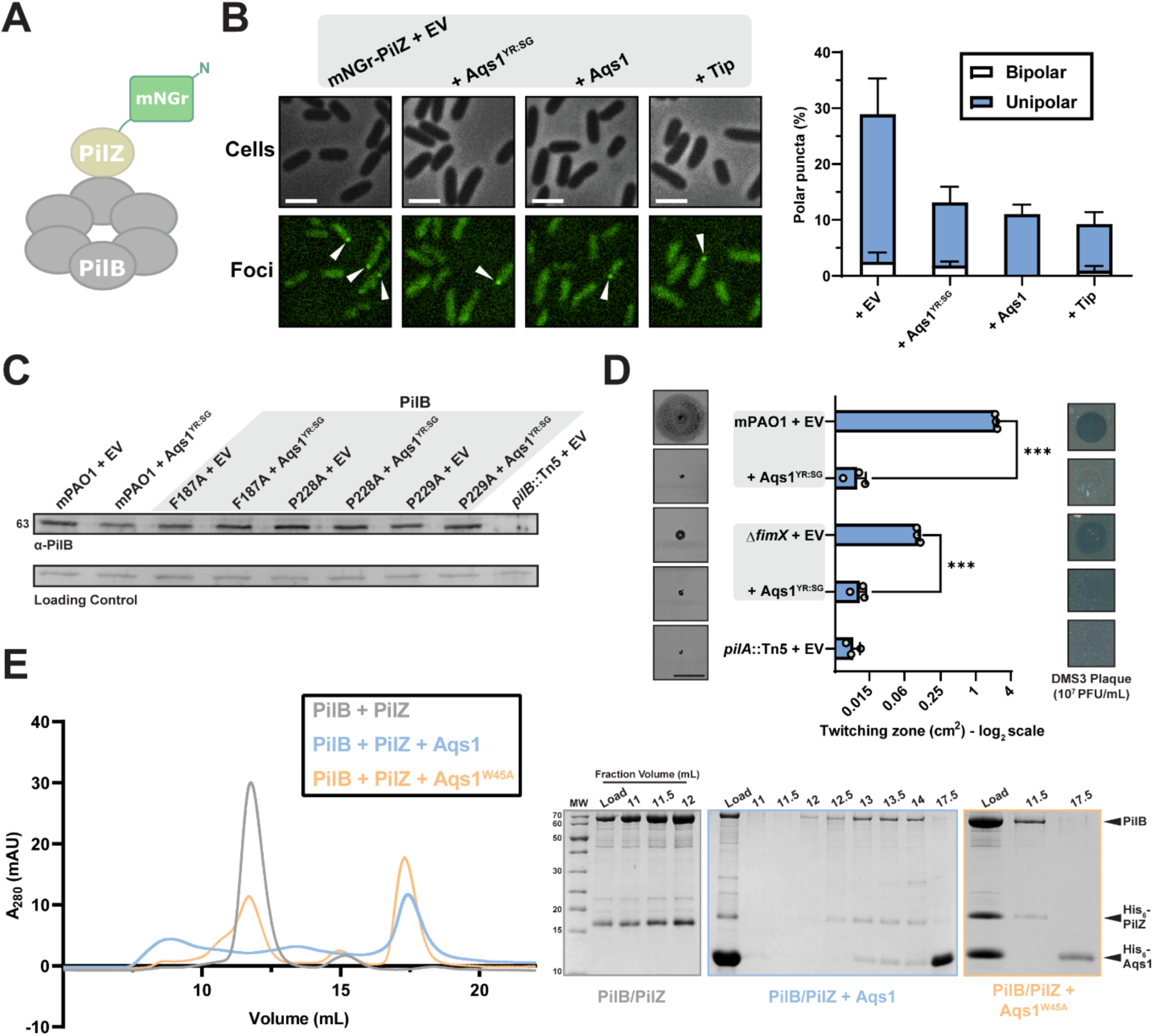
Aqs1 inhibits PilB by disrupting oligomerization. **(A)** Cartoon representation of N-terminal mNeonGreen PilZ (mNGr-PilZ) proxy. **(B)** Representative images of cells expressing mNGr-PilZ with background fluorescence subtracted. Strains grouped in the grey box indicate the same background. All scale bars = 2 µm. Quantification of cells scored for bipolar or unipolar puncta and expressed as a percentage of total cells is shown to the right. Bars represent the means ± SD of three independent experiments. **(C)** Representative PilB Western blot of strains expressing His_6_-Aqs1^YR:SG^. Sample loading was normalized to a non-specific band shown below. Molecular weight is indicated to the left. Blot is representative of three independent experiments. Indicated point mutations grouped in the grey box are all chromosomally introduced to PilB. **(D)** Quantification of sub-agar stab twitching motility zones of mPAO1 or a Δ*fimX* mutant expressing His_6_-Aqs1^YR:SG^. Representative twitching zones are shown to the left. X-axis depicts a log_2_ scale. Bars represent the means of triplicate samples from three independent experiments ± SD. All scale bars = 1 cm. Boxes to the right show representative DMS3 plaquing on a bacterial lawn of the same strain. Strains grouped in the grey boxes indicate the same background. **(E)** Representative size-exclusion chromatogram showing the elution profile of the full length PilB and His_6_-PilZ complex in the presence or absence of His_6_-Aqs1 or the non-PilB binding Aqs1^W45A^ mutant. SDS-PAGE of the corresponding elution fractions is shown to the right. **EV**: empty pHERD30T vector, **Aqs1:** N-terminally His_6_-tagged Aqs1 in pHERD30T vector, **Aqs1^YR:SG^**: N-terminally His_6_-tagged Aqs1 Y39S R40G in pHERD30T vector, **Tip:** N-terminally His_6_-tagged Tip in pHERD30T vector. **\*\*\***: 0.001 ≥ *p* (Two-tailed Welch’s *t-*test).

This finding suggested that rather than inhibiting ATPase activity, Aqs1 likely alters PilB structure, stability, or its functional associations with T4P components. We first tested whether Aqs1^YR:SG^ expression reduced PilB levels in whole cells via Western blotting. Expressing Aqs1^YR:SG^ in the presence of wild type PilB or the N2-domain mutants had no impact on steady state levels in the cell (**Fig 4C**). Next, we considered if Aqs1 could block the functional input of PilB protein regulatory effectors, such as PilZ described above, or the cyclic-di-GMP binding protein, FimX^40,41^. These effectors independently associate with PilB and are essential for its normal function^6,19,41^. FimX mutants retain some twitching, phage susceptibility, and produce short pili^22,42^. Despite the already substantial Δ*fimX* motility defect, twitching was further decreased and DMS3 could not infect when Aqs1^YR:SG^ was expressed in this background (**Fig 4D**). Taken together, these data suggest that Aqs1-dependent T4P inhibition occurs in the absence of FimX, and thus it is not the mechanistic target.

PilB hexamerization is essential for function, with individual subunits lacking the necessary architecture for substrate and T4P machine recognition^26–28,43^. We next assessed whether addition of Aqs1 to stable PilB hexamers could disrupt stability of the quaternary complex. We co-expressed and purified full-length PilB along with PilZ for N-terminal stabilization, separately from Aqs1 or the non-PilB binding Aqs1^W45A^ mutant (**Fig 1C**). When PilB-PilZ alone were passed over a size-exclusion chromatography column, the complex eluted in a single peak corresponding to ∼500 kDa, or roughly a 6:6 ratio of PilB/PilZ, with both PilB and PilZ detectable in the fractions (**Fig 4E**). When we added Aqs1, this peak disappeared and was replaced by a peak at an estimated MW of ∼168 kDa which corresponds to a 2:2:4 ratio of PilB/PilZ/Aqs1. PilB, PilZ, and Aqs1 were all detected in these fractions, confirming that Aqs1 does not prevent PilZ binding to PilB. As expected, the Aqs1^W45A^ mutant did not interact with the complex, and no dissociation was observed (**Fig 4E**, **S5**). These data suggest that Aqs1 can disrupt the oligomeric state of PilB-PilZ, thereby preventing it from acting at cell poles.

### Aqs1 may disrupt PilB linker-dependent function

We next sought to determine the structural basis of PilB oligomer disruption by Aqs1. Crystal structures of PilB hexamers demonstrate elongated two-fold symmetry that is considered to be the functional state of the enzyme^26–28^ (**Fig 5A**). This conformation results from differential substrate occupancy of the ATP-binding active sites resulting in nucleotide-bound (closed) or nucleotide-free (open) states^28^. We suspected that Aqs1 might disrupt the interface between subunits and/or interfere with N2-domain-dependent conformational changes in one or more of these states. We structurally aligned the PilB-Aqs1 bound monomer Alphafold3 model with the *G. metallireducens* PilB structure (PDB 5TSH) to capture Aqs1 positioning relative to subunits in all conformations (**Fig 5A**). Surprisingly, in each instance Aqs1 was positioned facing outward from the torus and we saw minimal steric clashing that might suggest the disruption of inter-subunit contacts (**Fig 5A**). Expressing an Aqs1 D5G mutant to remove the relevant side chain or truncating the N-terminus did not impact the inhibition of twitching motility, suggesting that the modelled clash is an artifact of the alignment (**Fig S6**). Left without a clear mechanistic conclusion, we considered that we were missing relevant structural details. Notably, PilB crystal structures lack density for the most distal N-terminal domain(s)^26–28^. How these domains associate with the rest of PilB is poorly understood, but we reasoned that the N-terminal elements might be perturbed by Aqs1 binding at the N2-domain. We generated a PilB truncation that lacked the N1-domain but retained the linker (PilB ΔT148; **Fig 5B**) in addition to our previous N-terminally truncated construct that lacks both the N1-domian and linker (PilB ΔD180; **Fig 3C**) to perform another Aqs1 co-purification in *E. coli*. Unexpectedly, we recovered more linker-intact PilB ΔT148 than linker-truncated ΔD180 following Ni^2+^-affinity chromatography (**Fig 5B**) suggesting this segment may indeed provide stabilization.

**Figure 5:**
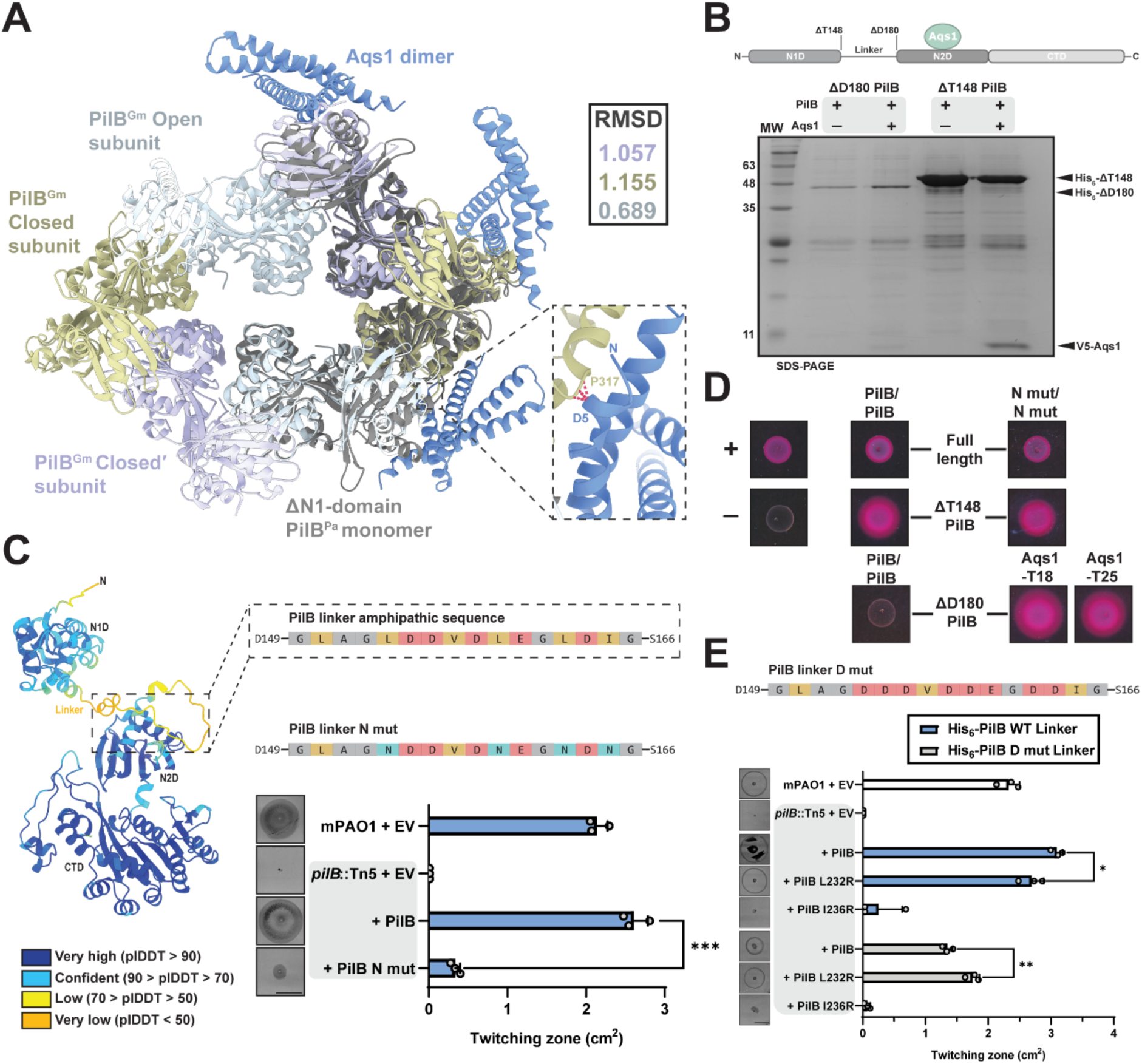
The PilB linker interacts with the N2-domain and is important for stabilization and function. **(A)** Alphafold3 model of the *P. aeruginosa* PilB monomer with N1-domain/linker hidden bound to the Aqs1 dimer structurally aligned onto the X-ray crystal structure of hexameric PilB from *G. metallireducens* (5TSH). The model is aligned onto each PilB open or closed subunit conformation. Inset depicts the single steric clash (red dashes, VDW radii overlap ≥ 0.6 Å) observed between Aqs1 and an adjacent PilB protomer. Each subunit pruned residue RMSD value is indicated in the top right. **(B)** SDS-PAGE of His_6_-PilB N-terminal truncations with or without V5-Aqs1 co-purification following Ni^2+^-affinity chromatography. Truncation boundaries are indicated in the cartoon above. Indicated PilB truncations are grouped in the grey boxes. **(C)** Alphafold3 model of the full length PilB monomer coloured by pLDDT score. The inset shows a portion of the PilB linker sequence highlighting hydrophobic amino acids. The PilB N mut sequence indicates where L154N, L159N, L162N, and I164N were introduced. Quantification of sub-agar stab twitching motility zones of mPAO1 *pilB*::Tn5 complemented with PilB or the PilB linker N mut shown below. Representative crystal violet-stained twitching zones are shown to the left. Strains grouped in the grey box indicate the same background. Bars represent the means of triplicate samples from three independent experiments ± SD. Expression was induced using 0.05% arabinose. **EV**: empty pHERD30T vector, **PilB:** PilB in pHERD30T vector, **PilB N mut:** PilB L154N, L159N, L162N, and I164N in pHERD30T vector. **(D)** Representative colonies showing self-interaction or pairwise interaction with Aqs1 (pink) between wildtype PilB or PilB N mut N-terminal truncations. Untagged T18/T25 plasmid and PilS-T18/PilS-T25 homodimers were used as negative and positive controls respectively. **(E)** Quantification of sub-agar stab twitching motility zones of mPAO1 *pilB*::Tn5 complemented with various PilB linker and/or N2-domain mutants. The PilB D mut sequence shows where L154D, L159D, and L162D were introduced. Representative twitching zones are shown to the left. Strains grouped in the grey box indicate the same background. Bars represent the means of duplicate samples from three independent experiments ± SD. All scale bars = 1 cm. **EV**: empty pHERD30T vector, **PilB WT Linker or indicated PilB point mutant:** N-terminally His_6_-tagged PilB or with an additional PilB point mutant in pHERD30T vector, **PilB D mut Linker or indicated PilB point mutant:** N-terminally His_6_-tagged PilB L154D, L159D, and L162D or with an additional PilB point mutant in pHERD30T vector. **\***: 0.05 ≥ *p* ≥ 0.01; **\*\***: 0.01 ≥ *p* ≥ 0.001; **\*\*\***: 0.001 ≥ *p* (Two-tailed Welch’s *t-*test).

This result was unanticipated as the linker lacks discernable density in crystal^26–28^ and cryo-EM^44^ structures of PilB homologues, and protein modelling tools struggle to define its structural features or estimate spatial positioning (**Fig 5C**, **S6**). The linker sequence is atypical for a potentially solvent-exposed segment, in that it comprises a cluster of hydrophobic amino acids interspaced by negatively charged and glycine residues (**Fig 5C**). We mutated several of these hydrophobic residues to asparagine (PilB N mut; **Fig 5C**), anticipating that if the linker stays free in the solvent as hypothesized^16,19^, these polar substitutions should be well tolerated. The PilB N mut construct complemented twitching poorly in a *pilB* mutant (**Fig 5C**) suggesting that hydrophobicity in the linker is important for PilB function. Some of these hydrophobic residues are separated by three residues and we hypothesized that the linker might adopt amphipathic helical secondary structure. However, introducing two adjacent helix-breaking proline residues into the sequence did not significantly decrease twitching (**Fig S6**). We next reasoned that the linker might be involved in PilB inter-subunit interaction. Both full-length PilB and the linker-intact PilB ΔT148 BACTH fusion constructs showed self-interaction while the N1-domain and no-linker PilB ΔD180 construct did not (**Fig 5D**). Of note, PilB ΔD180 continued to interact with Aqs1 under these conditions, indicating that the truncated protein is correctly expressed and folded. The PilB linker N mut truncations also demonstrated self-interaction under these conditions (**Fig 5D**). Together, these data suggest that this segment promotes PilB inter-subunit interaction but that linker hydrophobicity, while essential for PilB function, does not solely account for PilB-PilB contact.

From these data we considered that the linker might interact with the PilB N2-domain/C-terminal domain to promote oligomer stability (**Fig 5B**, **D**), and that Aqs1 could mechanistically block this feature to disrupt PilB oligomerization (**Fig 5E**). To evaluate this, we introduced positively charged arginine substitutions at L232 or I236 in the hydrophobic patch on the N2-domain (**Fig 2B**). L232R complemented twitching motility to near wild-type levels while I236R failed to restore motility (**Fig 5E**). In parallel, we mutated several of the hydrophobic residues in the linker to negatively charged aspartate (PilB D mut). Expressing this construct in a *pilB* mutant complemented twitching motility to only half of the control expressing wild-type PilB. However, twitching motility was significantly increased when the compensatory L232R positive charge was co-introduced in the N2-domain of the PilB D mut (**Fig 5E**). This suggests that the additional positive charge at the N2-domain compensates for the twitching defect imparted by increased negative charge in the linker. Together, these data imply that linker residues L154, L159, and L162 that are mutated in the PilB D mut normally contact residues like L232 in the N2-domain hydrophobic patch, which is also targeted by Aqs1, and that this interaction may support PilB complex stability.

## Discussion

Homologues of the PilB assembly ATPase are encoded by a variety of bacteria and deployed across systems with diverse functions^23^. Many of these contribute to virulence, enabling bacteria to adhere to and spread across surfaces supporting tissue invasion, uptake of exogenous DNA, protein toxin secretion, and biofilm formation^12,32,36,45–49^. Our finding that Aqs1, despite originating from a *P. aeruginosa* phage^2^, can disrupt T4P ATPase function in multiple bacteria is interesting as these proteins are not widely distributed beyond this genus. The Aqs proteins represent a unique class of phage-encoded PilB inhibitors with a mechanism of action distinct from other extension protein modulators^2,3,6,13,50,51^. Our data suggest that they exploit a conserved solvent-exposed hydrophobic patch on the N2-domain for PilB recognition (**Fig 2A**, **B**). This finding is consistent with previous work on Tip, where the authors confirmed through BACTH assays that it bound the N2-domain, but were unable to define the interaction at the sequence level^3^. We show that *P. aeruginosa* PilB P228, P229, and likely L232 side chains in this domain facilitate Aqs1 binding (**Fig 2C**, **3C**). Given the conservation of these critical residues between N2-domains of the Aqs1-binding ATPases (**Fig 2C**), Aqs1 is likely acting on the other PilB homologues tested by exploiting these same properties. By contrast, more divergent ATPases^23,28,52^ like TfpB^13^ lack the precise N2-domain sequence features that allow for Aqs1 interaction (**Fig S3**) and it fails to inhibit associated phenotypes (**Fig 1F**).

Once bound to the N2-domain, Aqs1 disrupts PilB polar retention in *P. aeruginosa* by disrupting hexamerization (**Fig 6A**). This might be achieved by displacing and/or preventing association of the PilB linker between the N1- and N2-domains (**Fig 5E).** Although it is a common feature of PilB homologues, the role of the linker is poorly understood due to its flexibility and thus lack of structural details^26–28^ (**Fig 5C**). From studies of various PilB homologues^19,53–55^, the N1-domain is thought to extend outward and away from the PilB torus to make contact with T4P structural components while the ATPase is docked at the base of the pilus nanomachine. Linker flexibility may therefore be needed to accommodate conformational rearrangements required for initial docking and/or to power pilus extension^28^. In *P. aeruginosa*, PilB does not engage with the T4P machine indefinitely, but rather is stochastically exchanged^35^ for dedicated retraction ATPases responsible for pilus depolymerization^45,56,57^. How then are the six N1-domains and flexible linkers stabilized during this transition? Our data indicate a functional interaction between the linker and a hydrophobic patch on the PilB N2-domain (**Fig 5E**), and this element in turn supports inter-subunit contact (**Fig 5D**). It is possible that while PilB is not docked with the pilus machinery, the linkers may withdraw to secure the N1-domains and thus stabilize the oligomer. If the linkers are unable to engage the N2-domain hydrophobic patches, such as when Aqs1 binds to the same site, this may be sufficient to weaken or block critical PilB inter-subunit interactions (**Fig 5D**). How the linker ultimately contacts the N2-domain, however, is unclear. Modelling the full length PilB hexamer with six PilZ proteins, to saturate each N1-domain, positions this subcomplex as engaged with the N2-/C-terminal domains (**Fig 6B**, **Fig S6**). Furthermore, the leucine side chains in the linker interact with the N2-domain hydrophobic patch from the same protomer in this model (**Fig 6B**). However, this prediction for the linker is low-confidence and the expected position error is high (**Fig 6B**, **S6**). As such, a separate scenario where this segment contacts the N2-domain of an adjacent protomer, or alternates between these states for hexamer stabilization, is also feasible (**Fig 6B**). Regardless, Aqs1 binding at the N2-domain would sterically disrupt these functional interactions, though more studies are needed to resolve PilB linker dynamics. Linker sequences of PilB homologues which bind to Aqs1 vary in length, amino acid composition, identity, and position of hydrophobic residues (**Fig 6C**). There is poor consensus between these sequences, suggesting that the precise linker and N2-domain engagement criteria may be species-specific. This supports that Aqs1 does not act on the linker directly but rather its broad-spectrum activity results from sterically decoupling these critical contacts with the more conserved N2-domain patch.

**Figure 6:**
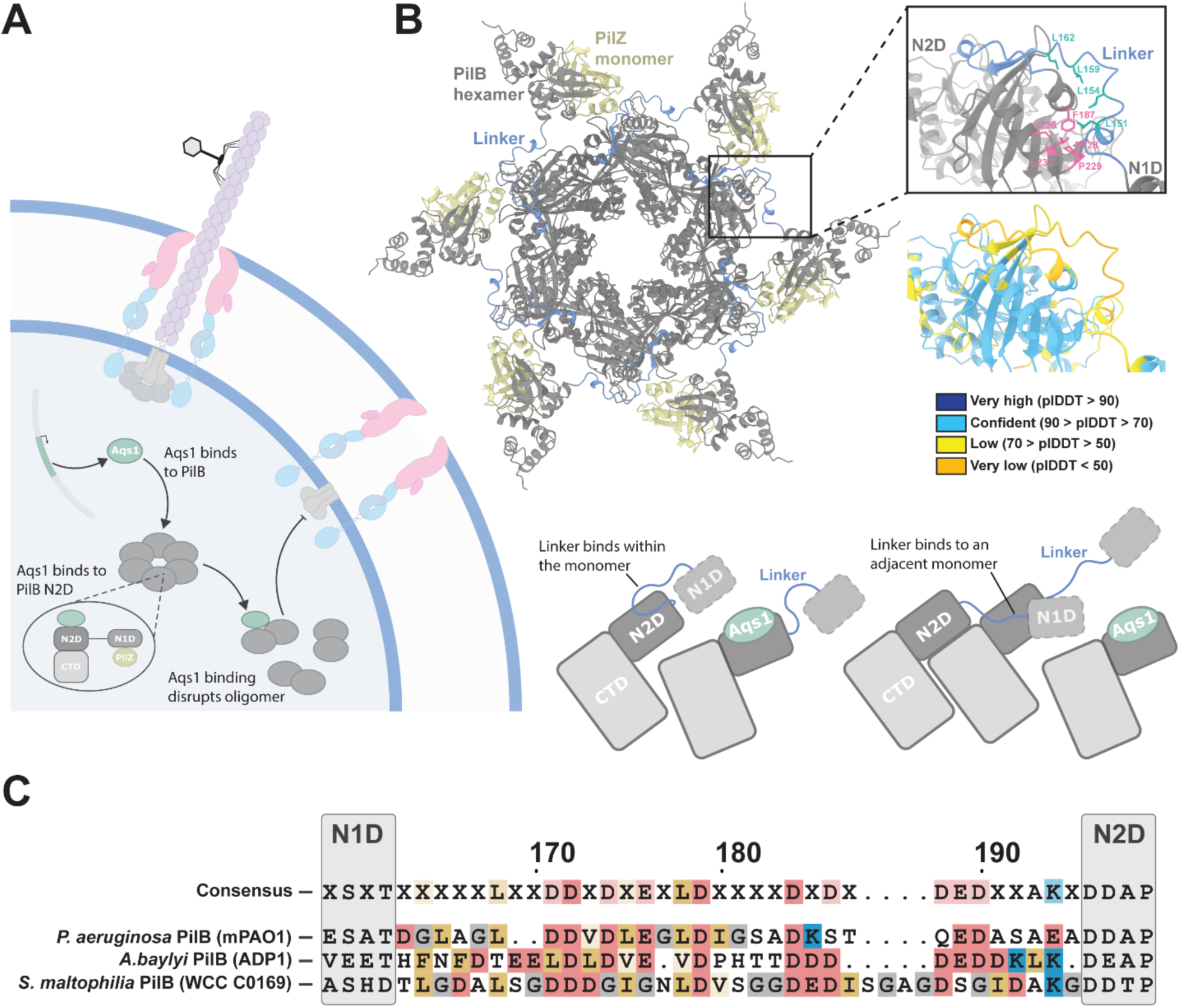
Mechanism of PilB inhibition by Aqs1. **(A)** To infect *P. aeruginosa*, phage DMS3 will first adsorb to T4P primary receptor. Once the phage has infected the cell it expresses Aqs1. Aqs1 targets the hexameric extension ATPase PilB and binds to the N2-domain (N2D). This binding will result in the disruption of PilB oligomerization to prevent engagement with pilus machines at the cell poles. **(B)** Alphafold3 model of the full length PilB hexamer (grey) with six PilZ monomers (tan). Each linker is indicated in blue. Inset shows the putative linker position with leucine side chains indicated in green relative to the N2-domain hydrophobic patch with relevant side chains indicated in pink. Below is the same inset coloured by pLDDT score. Aqs1 and the PilB flexible linker between the N1-domain (N1D) and N2-domain compete for an interaction surface on the PilB N2-domain. Linker binding to this allosteric site on the same or adjacent PilB subunit promotes hexamer stability, but this functional interaction is displaced by Aqs1. **(C)** Full length sequence alignment of Aqs1-binding PilB homologues highlighting the aligned linker sequences. Acidic, basic, branched hydrophobic/aromatic, and glycine residues are shaded in red, blue, tan, and grey respectively.

Our data emphasize the recognition by Aqs1 of an essential and conserved allosteric site unique to a subset of PilB homologues. This provides a template for the development of small molecules which mimic the Aqs1 mechanism of action, to act as broad-spectrum PilB inhibitors. Such compounds could serve to simultaneously limit the virulence of priority pathogens like *P. aeruginosa*, *S. maltophilia*, and possibly other high-risk organisms like *Acinetobacter baumannii*. By targeting the Aqs1-binding N2-domain allosteric region, such molecules may also avoid many of the toxicity challenges associated with traditional active-site inhibitors of ATPases^58^.

## Materials and Methods

### Bacterial strains and growth conditions

All strains and plasmids used in this study are outlined in Table S1. Bacterial strains were grown aerobically at 37 °C in Lennox lysogeny broth (LB) media (10 g/L tryptone, 5 g/L yeast extract, and 5 g/L NaCl), supplemented with the appropriate concentration of antibiotics where necessary unless otherwise specified. Antibiotics were used at the following concentrations: 15 µg/mL, 30 µg/mL, or 200 µg/mL of gentamicin for *E. coli*, *P. aeruginosa*, and *S. maltophilia* respectively unless otherwise indicated. 50 µg/mL of kanamycin. 100 µg/mL of ampicillin. 34 µg/mL of streptomycin. 34 µg/mL of chloramphenicol. Unless otherwise specified, arabinose was not supplemented into the media and experiments were performed with basal level gene expression in the absence of inducer. Plasmid constructs were generated using *E. coli* DH5α before introduction into the appropriate cellular background by heat shock (*E. coli*) or electroporation (*P. aeruginosa, E. coli*, or *S. maltophilia*) where appropriate.

*A. baylyi* cultures were grown at 30 °C in Miller LB medium and on agar supplemented with kanamycin (50 µg/ml), spectinomycin (60 µg/ml), and gentamycin (30 µg/ml) where appropriate.

### Plasmid and strain construction

Fragments of interest were PCR amplified from isolated genomic DNA apart from the mNGr-PilZ knock-in construct, MshE, and Aqs1^YR:SG^ ^W45A^ sequences which were instead synthesized by Genscript FLASH Gene synthesis. All PCR primers are listed in Table S2. For point mutations, two separate PCR amplified DNA fragments were stitched together through a second single overlap extension PCR, with each DNA overhang including the mutation of interest introduced during primer synthesis. For plasmid construction, the in-frame genetic insert and corresponding vector were digested with the appropriate FastDigest restriction enzymes (ThermoFisher Scientific) as per manufacturer instruction and ligated with T4 DNA ligase. Ligated products were transformed via heat shock into *E. coli* DH5α and selected on 1.5% agar LB media plates supplemented with the appropriate antibiotic. Following selection, single colonies were harvested, and the plasmid was mini prepped using a GeneJet plasmid prep kit (ThermoFisher Scientific) for sequence confirmation (Plasmidsaurus or Flow Genomics).

Genetic manipulation of mPAO1 chromosomal DNA was performed by double homologous recombination as previously described^22^. In brief, for genetic deletions, ∼500 bp upstream and downstream of the desired target sequence were PCR amplified and purified from isolated PAO1 genomic DNA. Upstream and downstream sequences were engineered to contain ∼50-100 bases of the 5ʹ or 3ʹ ends of the gene deletion target to ensure minimal disruption of adjacent genes. Upstream and downstream fragments were digested with the appropriate restriction enzymes along with the suicide vector backbone pEX18Gm. Digested fragments were ligated using T4 ligase and transformed into via heat shock into *E. coli* DH5α and selected for on 1.5% agar LB media plates supplemented with gentamicin. Purified and sequenced plasmids were then transformed into *E. coli* SM10 (λ*pir*). A single colony of SM10 harbouring the plasmid was grown overnight in 3 mL of LB media supplemented with gentamicin. The following day, the SM10 culture was mixed with the appropriate strain of mPAO1 overnight culture in a 1:1 ratio (by volume), pelleted by centrifugation, the resulting cell pellet was resuspended in 50 µL of LB and then spotted on 1.5% agar LB plates dried in a biological safety cabinet for 20-30 min. The spot was allowed to dry at room temperature before being incubated at 37 °C overnight. The following day, the cell spot was scraped into fresh LB media and plated onto premixed *Pseudomonas* isolation agar (Difco) supplemented with 100 µg/mL of gentamicin. The plates were incubated again at 37 °C overnight. Single colonies were then re-streaked again on *Pseudomonas* isolation agar plates supplemented with 100 µg/mL of gentamicin and incubated overnight at 37 °C. The next day, cells were streaked in a zig-zag pattern on 10% LB (1 g/L tryptone, 0.5 g/L yeast extract) with no added NaCl, 1.5% agar, and 5% sucrose media plates and incubated overnight at 37 °C. Single colonies from the sucrose media selection plate were patched again onto a fresh sucrose media plate as well as an LB media plate supplemented with gentamicin. Patches which grew on sucrose media but not LB with gentamicin were assessed for chromosomal deletion via colony PCR.

For chromosomal knock-in of a point mutation, the same protocol as outlined above was performed only differing at initial construct design. Knock-in constructs were instead generated ∼500 bp upstream or downstream of the mutation site through single overlap extension PCR. Overlap extension primers were also designed to silently introduce or remove a restriction cut site in the genome where applicable to facilitate downstream quality assurance. The remaining procedure was then carried out as described.

Mutants in *A. baylyi* were made using natural transformation as described previously^59^. Briefly, mutant constructs were made by splicing-by-overlap (SOE) PCR to stich (1) ∼1-3 kb of the homologous region upstream of the gene of interest, (2) the mutation where appropriate (for deletion by allelic replacement with an AbR cassette), and (3) ∼3 kb of the homologous downstream region. For the Δ*tfpB*::*gent^R^*strain CE986, the upstream region was amplified using F1 + R1 primers, and the downstream region was amplified using F2 + R2 primers, and the Gent^R^ AbR cassette was amplified with ABD123 (ATTCCGGGGATCCGTCGAC) and ABD124 (TGTAGGCTGGAGCTGCTTC). ∼20 bp of homology was built into the R1 and F2 primers to overlap with the AbR cassette insert, and a splicing by overlap extension (SOE) PCR reaction was performed using a mixture of the upstream and downstream regions and the AbR cassette middle region using F1 + R2 primers. SOE PCR products were added with 50 µL of overnight-grown culture to 450 µL of LB in 2-mL round-bottom microcentrifuge tubes (USA Scientific) and grown at 30 °C rotating on a roller drum for 3–5 h. Transformants were serially diluted and plated on LB and LB + antibiotic and then screened by PCR. To introduce mutations into different strain backgrounds, the mutated region was amplified from a template mutant strain using F1 + R2 primers and transformed into the appropriate strain background exactly as described above. The Δ*<u>tfpB</u>* deletion was transformed into CE2902 to create CE2908, and the Δ*vanAB::kan^R^*, P*_JExD_*-*his-aqs1* expression construct was transformed into CE22 to create CE2922. All strains were confirmed by PCR and/or nanopore sequencing (Plasmidsaurus).

The Aqs1 expression strain was constructed by placing His-Aqs1 under a crystal violet inducible P_JExD_- promoter^60^ at the *vanAB* locus as described previously with some modifications. First, we generated a Δ*vanAB::kan^R^* as proof of concept that crystal violet could induce higher expression by amplifying the *vanAB* locus containing the KanR cassette from a previously published ^13^ complementation strain using primers 317 + 1915 for the up arm, and 2142 + 176 from a previously published^61^ mRuby3 expression strain to amplify the down arm containing a ribosome binding site (RBS) and *mRuby3* at the same locus for the down arm. The P*_JExD_*-promoter region was amplified from plasmid pJC580. The SOE containing all three DNA pieces was amplified using the F1 + R2 primers and transformed into *A. baylyi* as described above and confirmed by sequencing. Induction of this strain with 250 nM crystal violet resulted in an increase in mRuby3 fluorescence, and this strain was then used as a template to generate the His-Aqs1 expression strain CE2902 by using primers 317 + 2140 to amplify the up arm containing the Δ*vanAB::kan^R^*, P*_JExD_*-, primers 406 + 176 to amplify the down arm at the same locus, and primers 4471 + 4468 to amplify *his-aqs1* from pHERD30T-*his*-*aqs1*^2^. The SOE PCR was amplified and transformed into *A. baylyi* exactly as described above and confirmed by nanopore sequencing.

### Pyocyanin production assays

Bacterial strains were inoculated into 3 mL of LB media liquid cultures supplemented with gentamicin and grown overnight at 37 °C. The next morning, 5 µL of overnight culture was sub-cultured into 3 mL of fresh LB media supplemented with gentamicin and grown at 37 °C to mid-logarithmic phase (OD_600_ ∼0.5-1.0). Cultures were OD_600_ normalized to 0.1 and 10 µL of cell suspension was spotted onto premixed *Pseudomonas* isolation agar (Difco) supplemented with gentamicin dried for 20-30 min is a biological safety cabinet. The spot was allowed to dry at room temperature before incubation overnight at 37 °C in a standing incubator. The following morning, plates were imaged with a scanner.

### Twitching motility assays

Bacterial strains were streaked onto 1.5% agar LB media plates supplemented with gentamicin where applicable and grown overnight at 37 °C. The following day, cells were picked using a sterile pipette tip and stabbed down to the bottom layer plastic of 1% agar LB media tissue culture treated plates, supplemented with gentamicin and arabinose where indicated, dried for 30 min in a biological safety cabinet. Plates were incubated overnight at 37 °C. The following day, the agar was carefully removed with a pipette tip and discarded. The twitching zones adhered to the plate were stained with 0.1% crystal violet dye for a minimum of 10 min, washed twice with DI water, and left to air dry overnight at room temperature before imaging with a scanner. Twitching zone areas were then digitally traced using ImageJ, and either duplicates or triplicates were averaged.

### Phage-susceptibility assays

Bacterial strains were inoculated into 3 mL of LB media liquid cultures supplemented with gentamicin and grown overnight at 37 °C. The next morning, 5 µL of overnight culture was sub-cultured into 3 mL of fresh LB media supplemented with gentamicin and grown at 37 °C to mid-logarithmic phase (OD_600_ ∼0.5-1.0). For plate-based assays, 20 µL of OD_600_ normalized bacterial suspension was mixed with 10 mL of 1% agar LB media supplemented with gentamicin and 0.1% arabinose then poured into a 60 mm round plate. The plate was dried in a biological safety cabinet for 30 min before spotting 10 µL of 10^7^ PFU/mL of DMS3 stock on top. The plate was incubated overnight at room temperature. The following day, the plate was incubated again at 37 °C for 2-3 hours before imaging with a scanner.

For liquid-based assays, cultures were OD_600_ normalized to 0.1 and 2 µL of cell suspension was added to 200 µL of LB media supplemented with gentamicin total in a flat bottom 96 well plate. To each well, 1 µL of 10^5^ PFU/mL DMS3 phage stock was also added for a final MOI of ∼0.001. The plate lid was sealed with masking tape and then incubated overnight in a 37 °C shaking plate reader, with OD_600_ readings taken every 30 min for 16-18 hours. Curves were normalized to LB media blank wells in the same plate.

### Bacterial 2-hybrid (BACTH) assay

Single colonies of *E. coli* BTH101 co-transformed with both T18/T25 tagged target proteins were inoculated into 3 mL LB media overnight cultures supplemented with kanamycin, ampicillin, and 0.5 mM IPTG. The following morning, the cultures were normalized to an OD_600_ of 0.1 in fresh LB and 4 µL was spotted onto MacConkey agar plates (40 g/L, Difco) supplemented with kanamycin, ampicillin, 0.5 mM IPTG, and 1% maltose pre-dried for 40 min in a biological safety cabinet. The spots were allowed to dry at room temperature before incubation at 30 °C in a standing incubator. The plates were imaged after 24 hours. For each assay, untagged T18/T25 plasmid and PilS-T18/PilS-T25 homodimers^62^ were used as negative and positive controls respectively.

### Bacterial growth curves

Bacterial strains were inoculated into 3 mL of LB media liquid cultures supplemented with gentamicin and grown overnight at 37 °C. The next morning, 5 µL of overnight culture was sub-cultured into 3 mL of fresh LB media supplemented with gentamicin and grown at 37 °C to mid-logarithmic phase (OD_600_ ∼0.5-1.0). Cultures were OD_600_ normalized to 0.1 and 2 µL of cell suspension was added to 200 µL of LB media total supplemented with gentamicin and arabinose where indicated in a flat bottom 96 well plate. The plate lid was sealed with masking tape and then incubated overnight in a 37 °C shaking plate reader, with OD_600_ readings taken every 30 min for 16-18 hours. Curves were normalized to LB media blank wells in the same plate.

### Protein modelling and sequence alignments

Alphafold3 was used to predict protein structures, complexes, and generate predicted aligned error plots using the unmodified default settings^38^. All models and plots were visualized in UCSF ChimeraX^63^. All sequences were aligned using Geneious Prime and visualized using ESPript3^64^.

### Small scale co-purification and pull-down experiments

Each N-terminally truncated PilB variant, linker intact PilB (M1-D180 and M1-T148 respectively), and/or Aqs1 were subcloned into pETDuet-1 in the first site with an N-terminal His_6_-tag or the second site with an N-terminal V5 epitope tag respectively for inducible expression. PilB was purified as previously described with some alterations^22^. Briefly, ΔN1D-PilB expressing overnight cultures of *E. coli* BL21-pLysS (DE3) were sub-cultured (1:100) in 50 mL of LB supplemented with ampicillin and incubated at 37°C with shaking until mid-logarithmic phase (∼0.4-0.8 OD_600_). Cells were induced with 0.5 mM IPTG overnight at 18°C with shaking. The next morning, cells were harvested by centrifugation at 3500 xg for 20 min. Cell pellets were resuspended in lysis buffer (50 mM HEPES pH 7.3, 250 mM NaCl, 10 mM imidazole) and incubated on ice with 20mg of lysozyme for 15 min with periodic inversion. Cells were then lysed using sonication and the supernatant clarified through centrifugation at 31000 xg for 30 min. At this stage an input sample was taken for each condition at which point the remaining supernatant was incubated with 700 µL of Ni^2+^-NTA resin (Millipore) pre-equilibrated with lysis-buffer for 1 hour on ice with gentle shaking. The resin was washed with 10 mL of equilibration containing 30 mM imidazole buffer before elution with 1 mL of equilibration buffer containing 250 mM imidazole.

The elution fractions were then separated on a 15% SDS-polyacrylamide gel. Samples were transferred to a nitrocellulose membrane at 225 mA for 1 hour. The membranes were blocked with 5% (w/v) skim milk (Bioshop) dissolved in 1X Tris-buffered saline (TBS: 20 mM Tris-HCl pH 7.6, 150 mM NaCl) for 2 hours. Primary incubation was performed using a 1:5000 dilution for both anti-His and anti-V5 (Genscript and Sigma respectively). The membranes were washed 4 times for 5 min with 1X TBS before incubation using alkaline phosphatase-conjugated goat-anti rabbit and goat-anti mouse IgG secondary antibody (BioRad) in a 1:3500 dilution. The membranes were washed 4 times with 1X TBS and developed using alkaline phosphatase developing reagent according to manufacturer’s protocol, before imaging the membranes in an Azure Biosystems 400 transilluminator.

### Western blot analysis of whole cell lysates

Whole cell lysates were first separated on 12% SDS-polyacrylamide gels. Samples were transferred to a nitrocellulose membrane at 225 mA for 1 hour. The membranes were blocked with 5% (w/v) skim milk (Bioshop) dissolved in 1X Tris-buffered saline (TBS: 20 mM Tris-HCl pH 7.6, 150 mM NaCl) for 2 hours. Primary incubation was performed as previously described using rabbit antisera^22^. The membranes were washed 4 times for 5 min with 1X TBS before incubation using alkaline phosphatase-conjugated goat-anti rabbit IgG secondary antibody (BioRad) in a 1:3500 dilution. The membranes were washed 4 times with 1X TBS and developed using alkaline phosphatase developing reagent according to manufacturer’s protocol, before imaging the membranes in an Azure Biosystems 400 transilluminator.

### Fluorescence microscopy experiments

Pilus labeling in P*. aeruginosa* was performed exactly as described previously^22^. 100 µl of overnight cultures of *P. aeruginosa* were spotted onto prewarmed LB agar plates and grown upright at 37 °C for 4-5 hours. Plate-grown cells were then scraped off of plates using a P200 pipette tip and gently resuspended into 100 µl of fresh LB. AlexaFluor488-maleimide (Sigma) was added to the cell suspension for a final concentration of 50 ng/µl and incubated for 45-60 min in the dark at room temperature. Cells were washed once with 200 µl fresh LB using centrifugation settings of 18,000 x*g* for 1 min. Cell were resuspended into 200 µl fresh LB. mNeonGreen-PilZ strains were prepared exactly the same way except with gentamycin supplemented in the medium and with no AlexaFluor488-maleimide labeling steps. All imaging was performed under 1% agarose pads made with PBS solution. Cell bodies were imaged using phase-contrast microscopy on a Nikon Ti2-E microscope using a Plan Apo 100X oil immersion objective, a GFP/FITC/Cy2 filter set for pili and mNeonGreen-PilZ, a Hamamatsu ORCA-Fusion Gen-III cCMOS camera, and Nikon NIS Elements Imaging Software.

Pilus labeling in *A. baylyi* was performed as described previously^59^. Briefly, 100 µl of overnight cultures grown with or without 250 nM crystal violet (CV) was added to 900 µl of fresh LB with or without 250 nM (only added to cultures grown overnight with CV) in a 1.5 ml microcentrifuge, and cells were grown at 30°C rotating on a roller drum for 70 min. Cells were then centrifuged at 18,000 × *g* for 1 min and resuspended in 50 µl of LB before labeling with 25 µg/ml Alexa-Fluor488 C5-maleimide (AF488-mal) (Thermo Fisher Scientific) for 15 min at room temperature. Labeled cells were centrifuged, washed three times with 100 µl of PBS and resuspended in 5-40 µl PBS. Cell bodies were imaged using phase-contrast microscopy, while labeled pili were imaged using fluorescence microscopy using the exact microscope set up described above. Cell numbers and the percentage of cells making pili over a 1 min time window with images captured at 10 s/frame were quantified manually using Fiji. All imaging was performed under 1% agarose pads made with PBS solution.

### Large scale protein purifications and size-exclusion chromatography

Full length PilB and PilZ were cloned into pCDF and pETDuet-1 in the first site with an N-terminal His_6_-tag leaving the second site empty respectively. Aqs1 or Aqs1^W45A^ were cloned into p15TVL with an N-terminal His_6_-tag as previously described^2^. Each protein was purified in the same manner. Briefly, overnight cultures of *E. coli* BL21 (DE3) co-transformed with PilB-pCDF and PilZ-pETDuet-1 or Aqs1/Aqs1^W45A^ alone were sub-cultured (1:100) in 1 L of LB supplemented with ampicillin and streptomycin or ampicillin respectively and incubated at 37°C with shaking until mid-logarithmic phase (∼0.4-0.8 OD_600_). Cells were induced with 1 mM IPTG overnight at 16°C with shaking. The next morning, cells were harvested by centrifugation at 3500 xg for 20 min. Cell pellets were resuspended in lysis buffer (50 mM Tris pH 8.0, 250 mM NaCl, 10 mM imidazole) and incubated on ice with 20 mg of lysozyme for 15 min with periodic inversion. Cells were then lysed using sonication and the supernatant clarified through centrifugation at 31000 xg for 30 min. Each supernatant was incubated with ∼1 mL of Ni^2+^-NTA resin (Millipore) pre-equilibrated with lysis-buffer for 1 hour on ice with gentle shaking. The resin was washed with 15 mL of equilibration buffer containing 30 mM imidazole before elution with 5 mL equilibration buffer containing 300 mM imidazole. The elution fractions were dialyzed overnight at 4°C into 50 mM Tris pH 8.0, 250 mM NaCl, and 1 mM of DTT. The next morning, each sample was concentrated using a 10 kDa cutoff Amicon concentrator to ∼700 µL. The proteins were loaded using 500 µL sample loop onto an S200 Increase 10/300 size exclusion chromatography column (Cytiva). Elution fractions containing PilB and PilZ or either Aqs1 variant were pooled and further concentrated as needed.

For the experimental runs, 500 µL of the PilB-PilZ complex was incubated with 200 µL of buffer or whichever Aqs1 variant at 3X the concentration. The protein samples were incubated at room temperature for 1 hour before loading onto an S200 Increase 10/300 size exclusion chromatography column using the 500 µL sample loop. Protein content in each elution fraction was validated using SDS-PAGE.

For co-expression of all PilB, PilZ, and Aqs1 together, Aqs1 and PilB were cloned into pETDuet-1 in the first site with an N-terminal His_6_-tag and untagged second site respectively. PilZ was cloned into the pACYDuet untagged second site. Both constructs were co-transformed into *E. coli* BL21 (DE3) for expression and purification as described above, supplementing the LB media with ampicillin and chloramphenicol, and without dialysis or size exclusion chromatography.

### Natural transformation assays

Transformation assays in *A. baylyi* were performed exactly as described previously^59^ with some differences. Briefly, strains were grown overnight in LB at 30°C on a roller drum with or without 250 nM crystal violet to induce Aqs1 expression. Then, 50 µl of overnight culture was subcultured into 450 µl of fresh LB medium with or without 250 nM of crystal violet (only added to the same cultures grown with it overnight), and 50 ng of transforming DNA (tDNA) (a ∼7 kb PCR product containing Δ*pilT*::*spec* amplified using primers CE49 + CE50) was used for all transformation assays. tDNA was quantified using a Qubit (Thermo Fisher Scientific) following standard Qubit protocols. Transformations were incubated with end-over-end rotation on a roller drum at 30°C for 5 h and then plated for quantification on LB + antibiotic plates (to quantify transformants) and on plain LB plates (to quantify total viable counts). Data are reported as the transformation frequency, which is defined as the (CFU/mL transformants)/(total CFU/mL).

### T2SS elastase secretion assays

*P. aeruginosa* strains were grown overnight in 3 mL LB media cultures supplemented with gentamicin and 0.1% arabinose. Each culture was then OD_600_ normalized to the lowest value with fresh LB media. 1 mL of each OD_600_ normalized culture was then centrifuged for 2 min at 21000 xg and the supernatant removed into a fresh 1.5 mL microcentrifuge tube. Approximately 80 mg of elastin-Congo red powder (Sigma) was then added to each tube. The exact mass of powder in each tube was recorded for later analysis. The tubes were vortexed vigorously to agitate the insoluble powder before then being incubated at 37 °C with shaking for 1, 3, and 5 hours. At each time point, the tubes were centrifuged for 2 min at 21000 xg to pellet the powder and 100 µL of supernatant was carefully removed into a 96-well plate to record absorbance at 495 nm. The tubes were vigorously vortexed again before returning to the shaking incubator. Final absorbance values were normalized to the exact mass of elastin-Congo red in each sample tube.

### Statistical Analyses

Microsoft Excel was used to determine averages of technical replicates and GraphPad Prism 8 was used to plot averages of biological replicates and calculate standard deviations. Natural transformation assay statistics were performed in Excel. Where indicated, an unpaired *t*-test with Welch’s correction, Sidak’s multiple comparisons test using log-transformed data, or a one-way analysis of variance (ANOVA) with Dunnett’s test for multiple comparison to a control strain was performed. Significance metrics are indicated in the corresponding figure captions.

## Acknowledgements

We thank Dr. P. Lynne Howell and Ian Yen for the PilB^Ab^, TfpB, and CpaF BACTH constructs as well as for helpful discussions and insights. We thank the Joint BioEnergy Institute for the Jungle Express promoter strain. We also thank Dr. Sara Andres, Caitlin Doubleday, and members of the Andres lab for use of equipment and their assistance in purifying some of the proteins used in this study. This work was supported by a Canadian Institutes of Health Research Project grant to LLB (PJT-156080) and to K.L.M. (PJT-203903). LLB is the Canada Research Chair in Microbe-Surface Interactions (CRC 2021–00103) and KLM is the Canada Research Chair in Bacteriophage Biology and Therapeutics (CRC-2023-00010). This work was also supported by a National Institutes of Health grant to CKE (R35GM150916). CKE is a Damon Runyon-Marilyn and Scott Urdang Breakthrough Scientist supported by the Damon Runyon Cancer Research Foundation (DFS6023). NR holds an NSERC CGS-D award.

