## Supplementary Figures and Tables for "A broad-spectrum phage-encoded mechanism to disarm bacterial type IV filaments"

**Supplemental information**

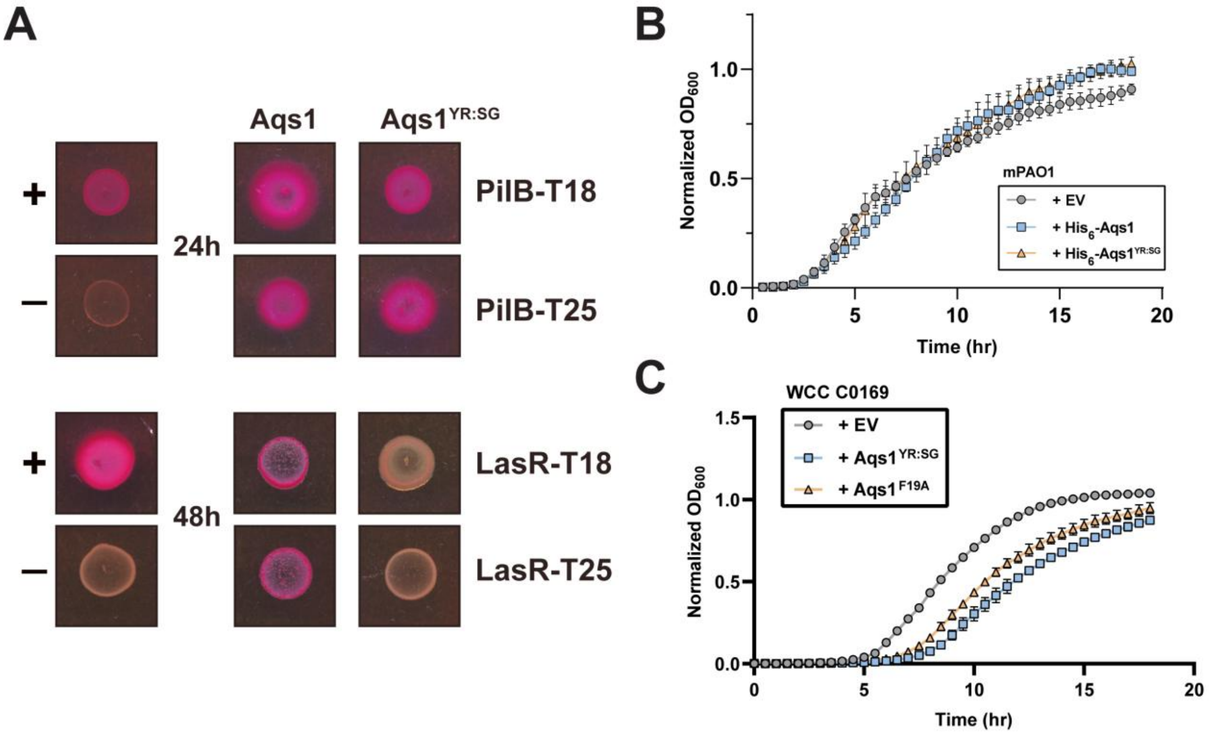


**Figure S1: Aqs1 and Aqs1$^{YR:SG}$ target specificity and effect on bacterial growth. (A)** Representative colonies showing pairwise interaction (pink) between Aqs1 variants with PilB or LasR following incubation at 24 and 48 hours respectively. Untagged T18/T25 plasmid and PilS-T18/PilS-T25 homodimers were used as negative and positive controls respectively. **(B)** Bacterial growth curves across 18 hours in LB media of mPAO1 expressing the indicated His$_6$-Aqs1 variant from pHERD30T (EV). Points represent the means of triplicate samples from three independent experiments ± SD. **(C)** Bacterial growth curves across 18 hours in LB media of WCC C0169 expressing the indicated Aqs1 variant from pBADGr (EV). Points represent the means of triplicate samples from three independent experiments ± SD. All Aqs1 expression was induced using 0.1% arabinose.

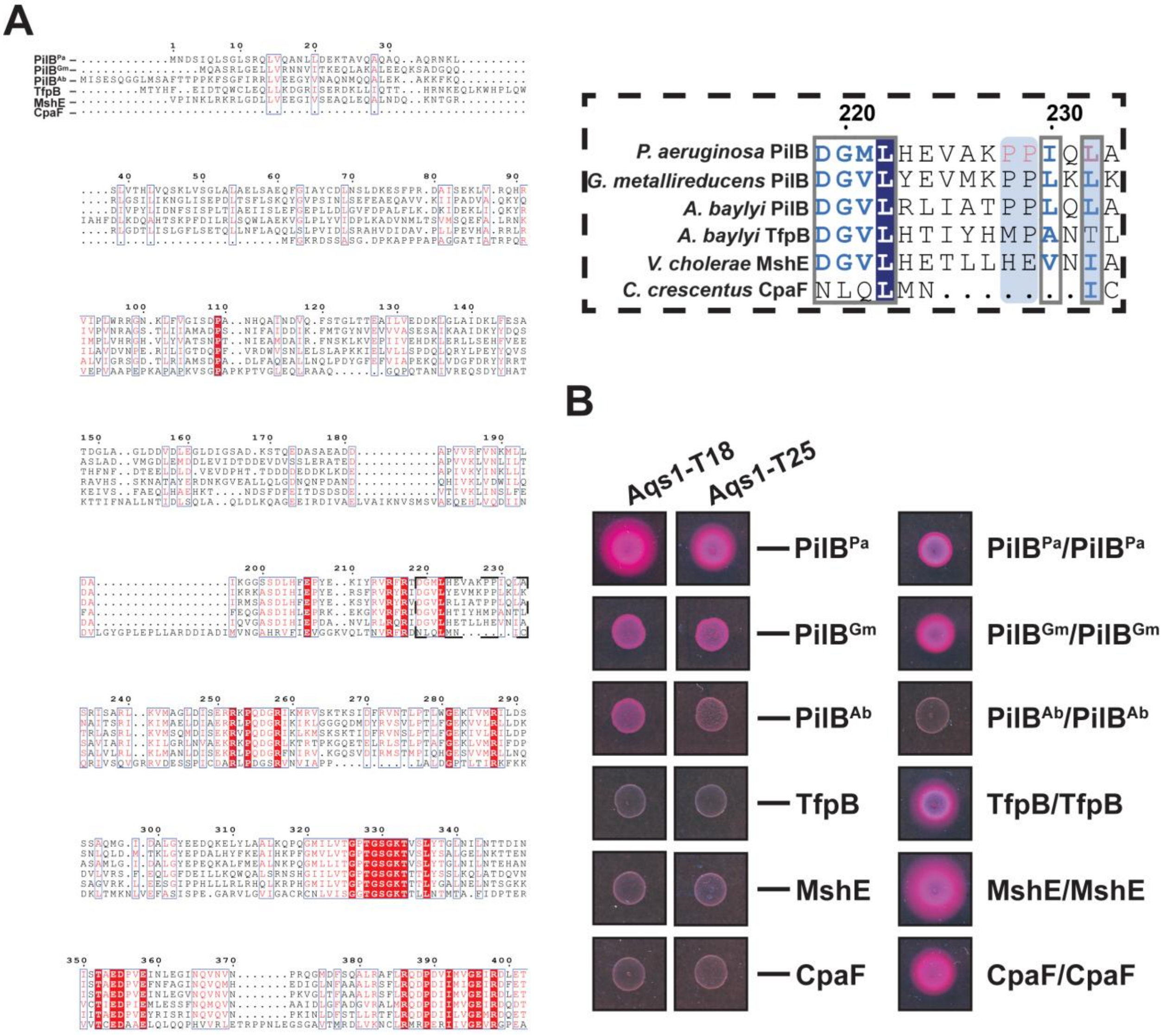


**Figure S2: The PilB monomer and Aqs1 dimer full model and T4 ATPase sequence alignment.** **(A)** Alphafold3 model of the full length PilB monomer and Aqs1 dimer coloured by pLDDT score. The PAE plot is shown below. **(B)** Translucent surface hydrophobicity overlayed onto the PilB N2-domain cartoon with relevant putative Aqs1-binding side chains highlighted in pink. **(C)** Full length sequence alignment of PilB homologues. Boxes indicate regions of sequence similarity. Residues shaded in red are identical across all sequences.

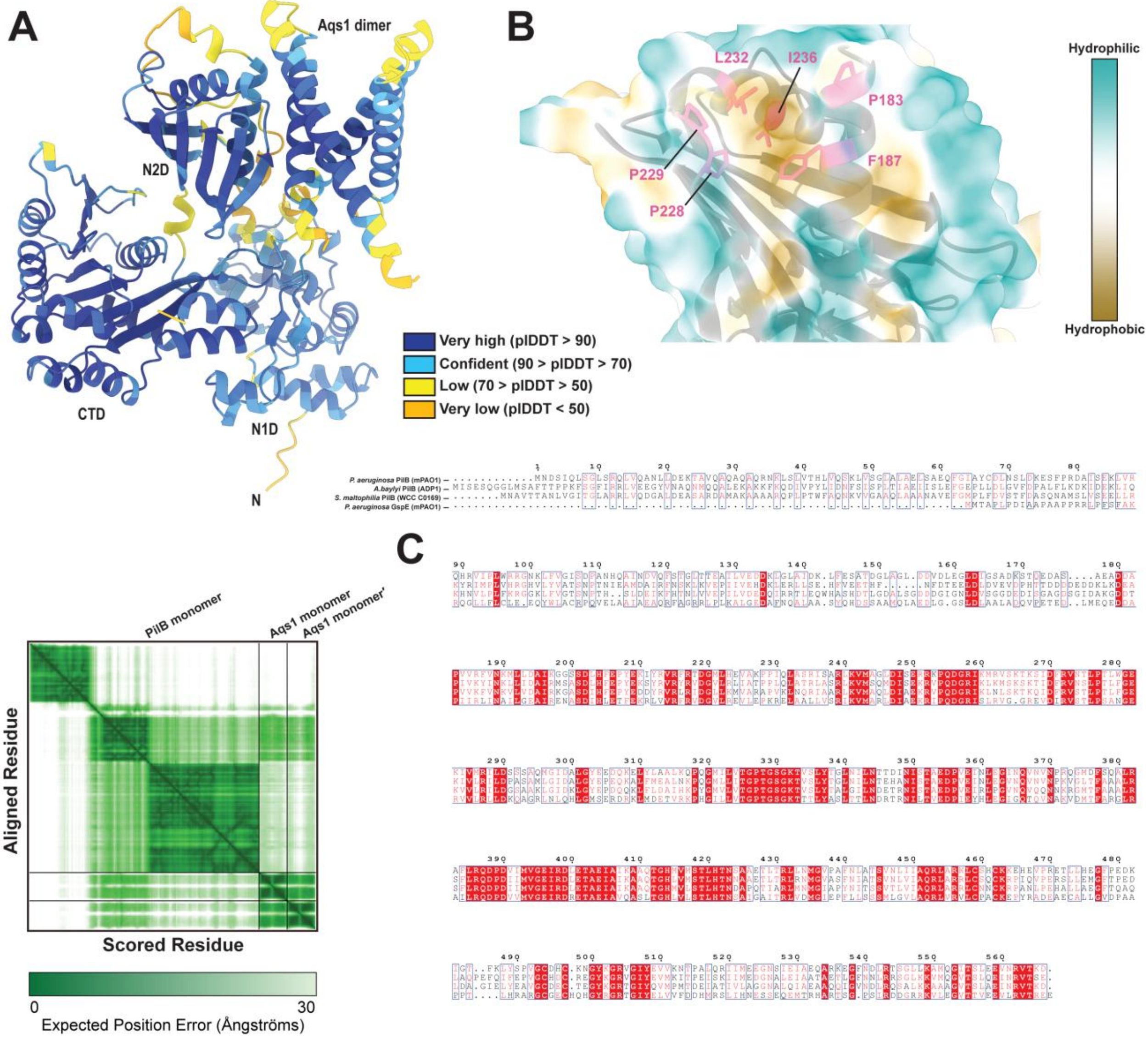


**Figure S3: T4 ATPases that lack certain sequence features do not interact with Aqs1. (A)** Full length sequence alignment of PilB homologues. Boxes indicate regions of sequence similarity. Residues shaded in red are identical across all sequences. Inset is an expanded region with relevant Aqs1-binding side chains from PilB$^{Pa}$ (P228, P229, and L232) shaded in pink. **(B)** Representative colonies showing pairwise interaction (pink) between PilB homologues from *Pseudomonas aeruginosa* (PilB$^{Pa}$), *Geobacter metallireducens* (PilB$^{Gm}$), *Acinetobacter baylyi* (PilB$^{Ab}$ and TfpB), *Vibrio cholerae* (MshE), and *Caulobacter crescentus* (CpaF) with Aqs1. ATPase self-interactions are also shown to the right.

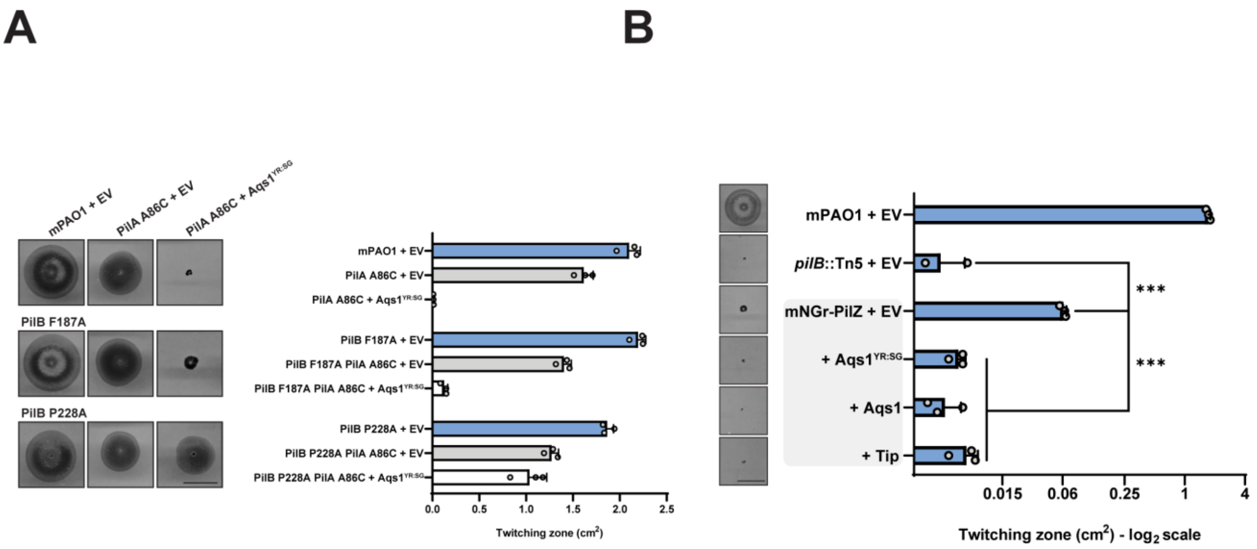


**Figure S4: All chromosomal mutants in this study retain twitching motility. (A)** Quantification of sub-agar stab twitching zone areas of PilA A86C mutants expressing Aqs1$^{YR:SG}$. Representative crystal violet stained twitching zones are shown to the left. Bars represent the means of triplicate samples from three independent experiments ± SD. **(B)** Quantification of sub-agar stab twitching zone areas of mNGr-PilZ mutant expressing Aqs protein homolgues. Representative twitching zones are shown to the left. Strains grouped in the grey box indicate the same background. Bars represent the means of triplicate samples from three independent experiments ± SD. All scale bars = 1 cm. **EV**: empty pHERD30T vector, **Aqs1$^{YR:SG}$**: N-terminally $His_6$-tagged Aqs1 Y39S R40G in pHERD30T vector, **Aqs1**: N-terminally $His_6$-tagged Aqs1 in pHERD30T vector, **Tip**: N-terminally $His_6$-tagged Tip in pHERD30T vector. *****: $0.001 \geq p$ (Two-tailed Welch's *t*-test).

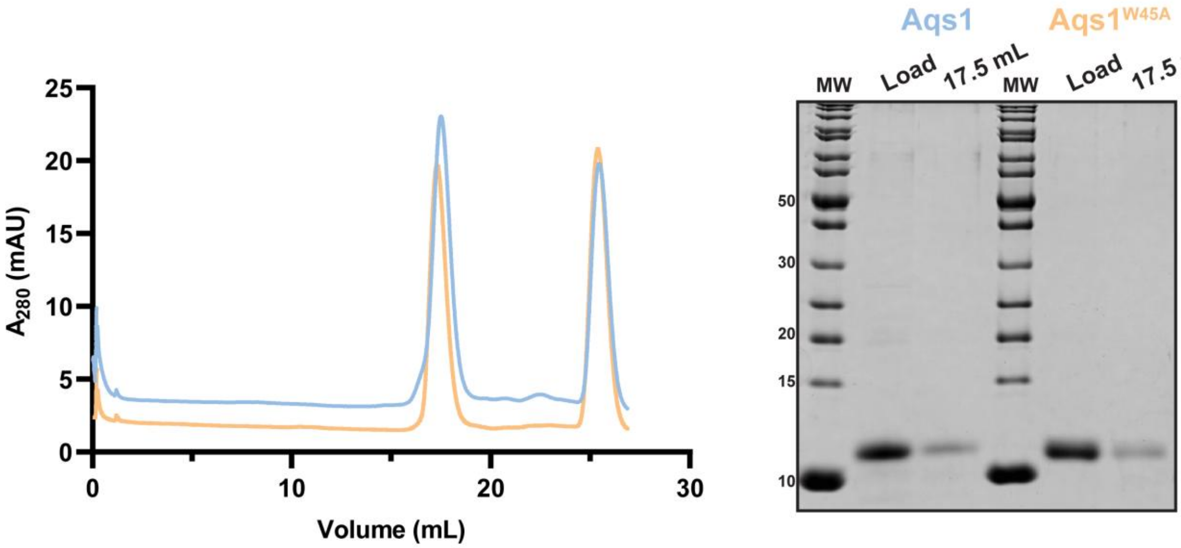


**Figure S5: Purification of His$_6$-Aqs1 or Aqs1$^{W45A}$ by size exclusion chromatography.** Size-exclusion chromatogram showing the elution profile of His$_6$-Aqs1 or His$_6$-Aqs1$^{W45A}$ alone. SDS-PAGE of the corresponding elution fractions is shown to the right.

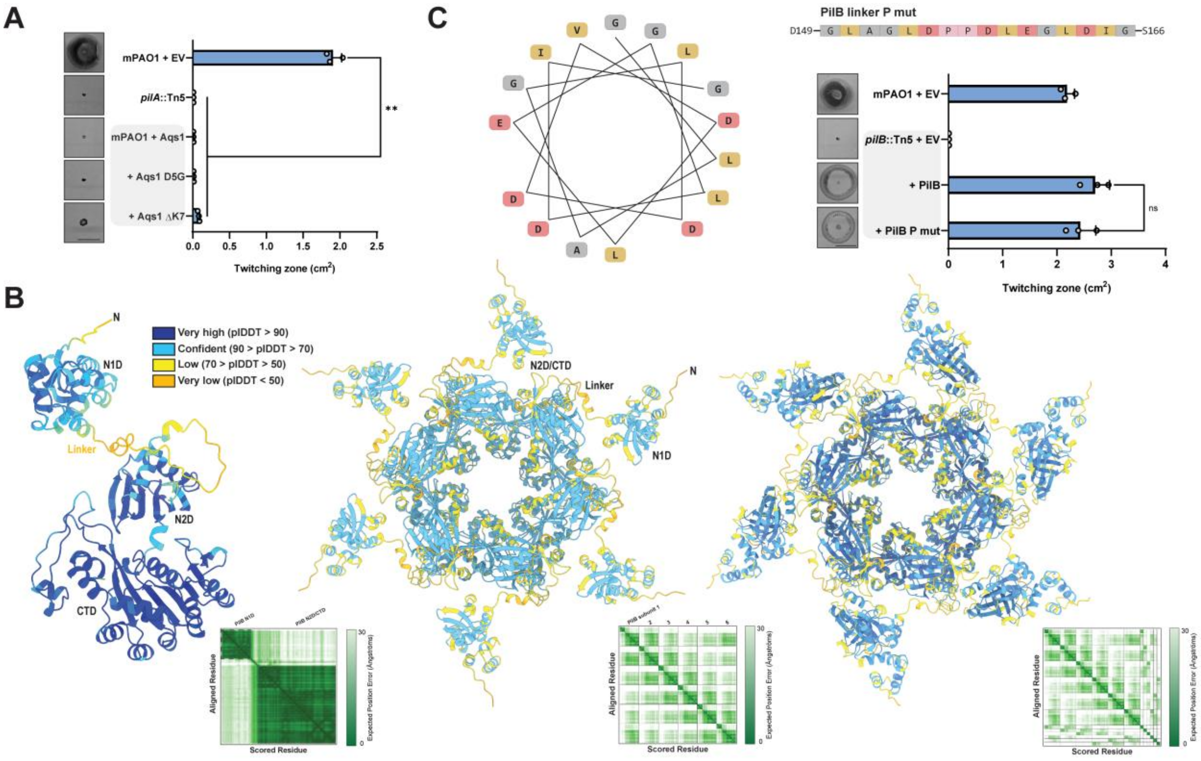


**Figure S6: The PilB linker has poorly defined structure or secondary structural elements.** **(A)** Quantification of sub-agar stab twitching zone areas of mPAO1 expressing Aqs1 N-terminal mutants. Representative crystal violet stained twitching zones are shown to the left. Bars represent the means of triplicate samples from three independent experiments ± SD. **(B)** PilB models coloured by pLDDT score. Full length PilB monomer**,** hexamer, and hexamer with six copies of PilZ generated using Alphafold3. Predicted aligned error (PAE) plots are shown below. **(C)** Helical wheel plot demonstrating a theoretical alpha-helical arrangement of residues in the PilB linker. Quantification of sub-agar stab twitching motility zones of mPAO1 *pilB*::Tn5 complemented with PilB or the PilB linker P mut shown above. Representative crystal violet stained twitching zones are shown to the left. Strains grouped in the grey box indicate the same background. Bars represent the means of triplicate samples from three independent experiments ± SD. All scale bars = 1 cm. **EV**: empty pHERD30T vector, **Aqs1:** N-terminally $His_6$-tagged Aqs1 in pHERD30T vector, **Aqs1 D5G:** N-terminally $His_6$-tagged Aqs1 D5G in pHERD30T vector, **Aqs1 ΔK7:** N-terminally $His_6$-tagged Aqs1 M1-L6 truncation in pHERD30T vector, **PilB:** PilB in pHERD30T vector, **PilB P mut:** PilB L156P and V157P in pHERD30T vector. **ns**: $p \geq 0.05$; ******: $0.01 \geq p \geq 0.001$ (Two-tailed Welch's *t*-test).

### Supplemental Tables

#### Table S1: All strains and plasmids used in this study.

| **Strains** | | |
|---|---|---|
| **Strain name** | **Genotype** | **Source** |
| ***E. coli* strains** | | |
| DH5α | *F-φ80lacZΔM15* Δ(*lacZYA-argF*)*U169 recA1 endA1 hsdR17*(*rk*−, *mk*+) *phoA supE44 thi-1 gyrA96 relA1 λ*− | Invitrogen |
| BL21 (DE3) | *fhuA2 [lon] ompT gal [dcm] ΔhsdS* | New England Biolabs |
| BL21 pLysS | *F–, ompT, hsdS*$_B$ *(r*$_B$*–, m*$_B$*–), dcm, gal, λ(DE3), pLysS, Cm*$^r$ | Novagen |
| BTH101 | Bacterial 2-hybrid strain | Euromedex |
| SM10 | *thi-1 thr leu tonA lacY supE recA*::*RP4-2-Tc*::*Mu* (*KmR*) | Invitrogen |
| PilS/PilS | BTH101 with PilS in pUT18C and pKT25 vectors | (1) |
| T18/T25 | BTH101 with empty pUT18C and pKT25 vectors | This study |
| PilB/PilB | BTH101 with PilB in pUT18C and pKT25 vectors | This study |
| PilB-T18/Aqs1-T25 | BTH101 with PilB in pUT18C and Aqs1 in pKT25 vectors | This study |
| PilB-T25/Aqs1-T18 | BTH101 with PilB in pKT25 and Aqs1 in pUT18C vectors | This study |
| PilB-T18/Aqs1 YR:SG -T25 | BTH101 with PilB in pUT18C and Aqs1 YR:SG in pKT25 vectors | This study |
| PilB-T25/Aqs1 YR:SG -T18 | BTH101 with PilB in pKT25 and Aqs1 YR:SG in pUT18C vectors | This study |
| LasR-T18/Aqs1-T25 | BTH101 with LasR in pUT18C and Aqs1 in pKT25 vectors | This study |
| LasR-T25/Aqs1-T18 | BTH101 with LasR in pKT25 and Aqs1 in pUT18C vectors | This study |
| LasR-T18/Aqs1 YR:SG -T25 | BTH101 with LasR in pUT18C and Aqs1 YR:SG in pKT25 vectors | This study |
| LasR-T25/Aqs1 YR:SG -T18 | BTH101 with LasR in pKT25 and Aqs1 YR:SG in pUT18C vectors | This study |
| PilB P183A-T18/Aqs1-T25 | BTH101 with PilB P183A in pUT18C and Aqs1 in pKT25 vectors | This study |
| PilB P183A-T25/Aqs1-T18 | BTH101 with PilB P183A in pKT25 and Aqs1 in pUT18C vectors | This study |
| PilB F187A-T18/Aqs1-T25 | BTH101 with PilB F187A in pUT18C and Aqs1 in pKT25 vectors | This study |
| PilB F187A-T25/Aqs1-T18 | BTH101 with PilB F187A in pKT25 and Aqs1 in pUT18C vectors | This study |
| PilB P228A-T18/Aqs1-T25 | BTH101 with PilB P228A in pUT18C and Aqs1 in pKT25 vectors | This study |
| PilB P228A-T25/Aqs1-T18 | BTH101 with PilB P228A in pKT25 and Aqs1 in pUT18C vectors | This study |
| PilB P229A-T18/Aqs1-T25 | BTH101 with PilB P229A in pUT18C and Aqs1 in pKT25 vectors | This study |

| | | |
|---|---|---|
| PilB P229A-T25/Aqs1-T18 | BTH101 with PilB P229A in pKT25 and Aqs1 in pUT18C vectors | This study |
| PilB L232A-T18/Aqs1-T25 | BTH101 with PilB L232A in pUT18C and Aqs1 in pKT25 vectors | This study |
| PilB L232A-T25/Aqs1-T18 | BTH101 with PilB L232A in pKT25 and Aqs1 in pUT18C vectors | This study |
| PilB I236A-T18/Aqs1-T25 | BTH101 with PilB I236A in pUT18C and Aqs1 in pKT25 vectors | This study |
| PilB I236A-T25/Aqs1-T18 | BTH101 with PilB I236A in pKT25 and Aqs1 in pUT18C vectors | This study |
| pETDuet-1 ΔN1D (ΔD180) PilB | BL21 pLysS with N-terminally truncated $His_6$-tagged PilB in the first site and empty second site for heterologous expression in *E. coli* | This study |
| pETDuet-1 Aqs1 | BL21 pLysS with an empty first site and V5-tagged Aqs1 in the second site for heterologous expression in *E. coli* | This study |
| pETDuet-1 ΔN1D PilB/Aqs1 | BL21 pLysS with N-terminally truncated $His_6$-tagged PilB in the first site and V5-tagged Aqs1 in the second site for heterologous expression in *E. coli* | This study |
| pETDuet-1 ΔN1D PilB F187A/Aqs1 | BL21 pLysS with N-terminally truncated $His_6$-tagged PilB F187A in the first site and V5-tagged Aqs1 in the second site for heterologous expression in *E. coli* | This study |
| pETDuet-1 ΔN1D PilB P228A/Aqs1 | BL21 pLysS with N-terminally truncated $His_6$-tagged PilB P228A in the first site and V5-tagged Aqs1 in the second site for heterologous expression in *E. coli* | This study |
| pETDuet-1 ΔN1D PilB P229A/Aqs1 | BL21 pLysS with N-terminally truncated $His_6$-tagged PilB P229A in the first site and V5-tagged Aqs1 in the second site for heterologous expression in *E. coli* | This study |
| PilB-T18/Aqs2-T25 | BTH101 with PilB in pUT18C and Aqs2 in pKT25 vectors | This study |
| PilB-T25/Aqs2-T18 | BTH101 with PilB in pKT25 and Aqs2 in pUT18C vectors | This study |
| PilB-T18/Aqs3-T25 | BTH101 with PilB in pUT18C and Aqs3 in pKT25 vectors | This study |
| PilB-T25/Aqs3-T18 | BTH101 with PilB in pKT25 and Aqs3 in pUT18C vectors | This study |
| PilB N1D-T18/Aqs1-T25 | BTH101 with the PilB N1D in pUT18C and Aqs2 in pKT25 vectors | This study |
| PilB N1D-T25/Aqs1-T18 | BTH101 with the PilB N1D in pKT25 and Aqs2 in pUT18C vectors | This study |
| PilB N2D/CTD-T18/Aqs1-T25 | BTH101 with N-terminally truncated PilB in pUT18C and Aqs2 in pKT25 vectors | This study |
| PilB N2D/CTD-T25/Aqs1-T18 | BTH101 with N-terminally truncated PilB in pKT25 and Aqs2 in pUT18C vectors | This study |
| PilB N1D-T18/PilZ-T25 | BTH101 with the PilB N1D in pUT18C and Aqs2 in pKT25 vectors | This study |
| PilB N1D-T25/PilZ-T18 | BTH101 with the PilB N1D in pKT25 and Aqs2 in pUT18C vectors | This study |
| PilB N2D/CTD-T18/PilZ-T25 | BTH101 with N-terminally truncated PilB in pUT18C and Aqs2 in pKT25 vectors | This study |
| PilB N2D/CTD-T25/PilZ-T18 | BTH101 with N-terminally truncated PilB in pKT25 and Aqs2 in pUT18C vectors | This study |
| PilB pCDF PilZ pETDuet-1 | BL21 with PilB in pCDF and $His_6$-PilZ in pETDuet-1 first site with empty second site for heterologous expression in *E. coli* | This study |
| Aqs1 p15TVL | BL21 with $His_6$-Aqs1 in p15TVL for heterologous expression in *E. coli* | (2) |

| | | |
|---|---|---|
| Aqs1 W45A p15TVL | BL21 with $His_6$-Aqs1 W45A in p15TVL for heterologous expression in *E. coli* | (2) |
| PilB+PilZ+Aqs1 | BL21 with $His_6$-Aqs1 and PilB in pETDuet-1 first and second sites respectively and PilZ in pACYDuet second site with empty first site | This study |
| pETDuet-1 ΔT148 PilB | BL21 pLysS with linker intact N-terminally truncated $His_6$-tagged PilB in the first site and empty second site for heterologous expression in *E. coli* | This study |
| pETDuet-1 ΔT148 PilB/Aqs1 | BL21 pLysS with linker intact N-terminally truncated $His_6$-tagged PilB in the first site and V5-tagged Aqs1 in the second site for heterologous expression in *E. coli* | This study |
| PilB ΔT148/PilB ΔT148 | BTH101 with PilB linker intact ΔT148 in pUT18C and pKT25 vectors | This study |
| PilB ΔT148 N mut/ PilB ΔT148 N mut | BTH101 with PilB linker intact ΔT148 in pUT18C and pKT25 vectors | This study |
| PilB N mut /PilB N mut | BTH101 with PilB L154N, L159N, L162N, and I164N in pUT18C and pKT25 vectors | This study |
| PilB ΔD180 N mut -T18/Aqs1-T25 | BTH101 with N-terminally truncated PilB in pUT18C and Aqs1 in pKT25 vectors | This study |
| PilB ΔD180 N mut-T25/Aqs1-T18 | BTH101 with N-terminally truncated PilB in pKT25 and Aqs1 in pUT18C vectors | This study |
| $PilB^{Gm}/PilB^{Gm}$ | BTH101 with PilB from *Geobacter metallireducens* in pUT18C and pKT25 vectors | This study |
| $PilB^{Gm}$-T18/Aqs1-T25 | BTH101 with PilB from *Geobacter metallireducens* in pUT18C and Aqs1 pKT25 vectors | This study |
| $PilB^{Gm}$-T25/Aqs1-T18 | BTH101 with PilB from *Geobacter metallireducens* in pKT25 and Aqs1 pUT18C vectors | This study |
| $PilB^{Ab}/PilB^{Ab}$ | BTH101 with PilB from *Acinetobacter baylyi* in pUT18C and pKT25 vectors | This study |
| $PilB^{Ab}$-T18/Aqs1-T25 | BTH101 with PilB from *Acinetobacter baylyi* in pUT18C and Aqs1 pKT25 vectors | This study |
| $PilB^{Ab}$-T25/Aqs1-T18 | BTH101 with PilB from *Acinetobacter baylyi* in pKT25 and Aqs1 pUT18C vectors | This study |
| TfpB/TfpB | BTH101 with TfpB from *Acinetobacter baylyi* in pUT18C and pKT25 vectors | This study |
| TfpB-T18/Aqs1-T25 | BTH101 with TfpB from *Acinetobacter baylyi* in pUT18C and Aqs1 pKT25 vectors | This study |
| TfpB-T25/Aqs1-T18 | BTH101 with TfpB from *Acinetobacter baylyi* in pKT25 and Aqs1 pUT18C vectors | This study |
| MshE/MshE | BTH101 with MshE from *Vibrio cholerae* in pUT18C and pKT25 vectors | This study |
| MshE-T18/Aqs1-T25 | BTH101 with MshE from *Vibrio cholerae* in pUT18C and Aqs1 pKT25 vectors | This study |
| MshE-T25/Aqs1-T18 | BTH101 with MshE from *Vibrio cholerae* in pKT25 and Aqs1 pUT18C vectors | This study |
| CpaF/ CpaF | BTH101 with CpaF from *Caulobacter crescentus* in pUT18 and pKNT25 vectors | This study |
| CpaF -T18/Aqs1-T25 | BTH101 with CpaF from *Caulobacter crescentus* in pUT18 and Aqs1 pKT25 vectors | This study |
| CpaF -T25/Aqs1-T18 | BTH101 with CpaF from *Caulobacter crescentus* in pKNT25 and Aqs1 pUT18C vectors | This study |
| pETDuet-1 ΔN1D $PilB^{Gm}$ | BL21 pLysS with linker intact N-terminally truncated $His_6$-tagged $PilB^{Gm}$ in the first site and empty second site for heterologous expression in *E. coli* | This study |

| | | |
|---|---|---|
| pETDuet-1 ΔN1D PilB$^{Gm}$/Aqs1 | BL21 pLysS with linker intact N-terminally truncated His$_6$-tagged PilB$^{Gm}$ in the first site and V5-tagged Aqs1 in the second site for heterologous expression in *E. coli* | This study |
| pETDuet-1 ΔN1D PilB$^{Ab}$ | BL21 pLysS with linker intact N-terminally truncated His$_6$-tagged PilB$^{Ab}$ in the first site and empty second site for heterologous expression in *E. coli* | This study |
| pETDuet-1 ΔN1D PilB$^{Ab}$/Aqs1 | BL21 pLysS with linker intact N-terminally truncated His$_6$-tagged PilB$^{Ab}$ in the first site and V5-tagged Aqs1 in the second site for heterologous expression in *E. coli* | This study |
| ***P. aeruginosa* strains** | | |
| mPAO1 | WT | (3) |
| *pilA*::Tn5 | mPAO1 *pilA* T5 transposon mutant | (3) |
| *pilB*::Tn5 | mPAO1 *pilB* T5 transposon mutant | (3) |
| Δ*fimX* | mPAO1 Δ*fimX* | (4) |
| Δ*fimX* + EV | mPAO1 Δ*fimX* + empty pHERD30T | This study |
| Δ*fimX* + Aqs1 YR:SG | mPAO1 Δ *fimX* + His$_6$-Aqs1 YR:SG pHERD30T | This study |
| Δ*pilZ* | mPAO1 Δ*pilZ* | (4) |
| mPAO1 + EV | WT *P. aeruginosa* with empty pHERD30T | (4) |
| mPAO1 + Aqs1 | WT *P. aeruginosa* with His$_6$-Aqs1 in pHERD30T | This study |
| mPAO1 + Aqs1 YR:SG | WT *P. aeruginosa* with His$_6$-Aqs1 YR:SG in pHERD30T | This study |
| mPAO1 + Aqs1 F19A | WT *P. aeruginosa* with His$_6$-Aqs1 F19A in pHERD30T | This study |
| mPAO1 + Aqs1 W45A | WT *P. aeruginosa* with His$_6$-Aqs1 W45A in pHERD30T | This study |
| *pilB*::Tn5 + EV | mPAO1 *pilB* T5 transposon mutant with empty pHERD30T | (4) |
| Δ*lasR* + EV | mPAO1 Δ*lasR* with empty pHERD30T | This study |
| *pilB*::Tn5 + PilB | mPAO1 *pilB* T5 transposon mutant with PilB in pHERD30T | (4) |
| *pilB*::Tn5 + PilB P183A | mPAO1 *pilB* T5 transposon mutant with PilB P183A in pHERD30T | This study |
| *pilB*::Tn5 + PilB F187A | mPAO1 *pilB* T5 transposon mutant with PilB F187A in pHERD30T | This study |
| *pilB*::Tn5 + PilB P228A | mPAO1 *pilB* T5 transposon mutant with PilB P228A in pHERD30T | This study |
| *pilB*::Tn5 + PilB P229A | mPAO1 *pilB* T5 transposon mutant with PilB P229A in pHERD30T | This study |
| *pilB*::Tn5 + PilB L232A | mPAO1 *pilB* T5 transposon mutant with PilB L232A in pHERD30T | This study |
| *pilB*::Tn5 + PilB I236A | mPAO1 *pilB* T5 transposon mutant with PilB I236A in pHERD30T | This study |
| PilB F187A + EV | mPAO1 with chromosomal F187A mutation with empty pHERD30T | This study |
| PilB P228A + EV | mPAO1 with chromosomal P228A mutation with empty pHERD30T | This study |

| | | |
|---|---|---|
| PilB P229A + EV | mPAO1 with chromosomal P229A mutation with empty pHERD30T | This study |
| PilB F187A + Aqs1 YR:SG | mPAO1 with chromosomal F187A mutation with $His_6$-Aqs1 YR:SG in pHERD30T | This study |
| PilB P228A + Aqs1 YR:SG | mPAO1 with chromosomal P228A mutation with $His_6$-Aqs1 YR:SG in pHERD30T | This study |
| PilB P229A + Aqs1 YR:SG | mPAO1 with chromosomal P229A mutation with $His_6$-Aqs1 YR:SG in pHERD30T | This study |
| PilA A86C + EV | mPAO1 PilA A86C with empty pHERD30T | This study |
| PilA A86C + Aqs1 YR:SG | mPAO1 PilA A86C with $His_6$-Aqs1 YR:SG in pHERD30T | This study |
| PilB F187A PilA A86C + EV | mPAO1 PilB F187A PilA A86C with empty pHERD30T | This study |
| PilB F187A PilA A86C + Aqs1 YR:SG | mPAO1 PilB F187A PilA A86C with $His_6$-Aqs1 YR:SG in pHERD30T | This study |
| PilB P228A PilA A86C + EV | mPAO1 PilB P228A PilA A86C with empty pHERD30T | This study |
| PilB P228A PilA A86C + Aqs1 YR:SG | mPAO1 PilB P228A PilA A86C with $His_6$-Aqs1 YR:SG in pHERD30T | This study |
| mPAO1 + Aqs2 | WT *P. aeruginosa* with $His_6$-Aqs2 in pHERD30T | This study |
| mPAO1 + Aqs3 | WT *P. aeruginosa* with $His_6$-Aqs3 in pHERD30T | This study |
| PilB P228A + Aqs1 | mPAO1 PilB P228A with $His_6$-Aqs1 in pHERD30T | This study |
| PilB P228A + Aqs2 | mPAO1 PilB P228A with $His_6$-Aqs2 in pHERD30T | This study |
| PilB P228A + Aqs3 | mPAO1 PilB P228A with $His_6$-Aqs3 in pHERD30T | This study |
| PilB P229A + Aqs1 | mPAO1 PilB P229A with $His_6$-Aqs1 in pHERD30T | This study |
| PilB P229A + Aqs2 | mPAO1 PilB P229A with $His_6$-Aqs2 in pHERD30T | This study |
| PilB P229A + Aqs3 | mPAO1 PilB P229A with $His_6$-Aqs3 in pHERD30T | This study |
| mNGr-PilZ + EV | mPAO1 with PilZ chromosomally tagged at the N-terminus with mNeonGreen with empty pHERD30T | This study |
| mNGr-PilZ + Aqs1 | mPAO1 with PilZ chromosomally tagged at the N-terminus with mNeonGreen and $His_6$-Aqs1 in pHERD30T | This study |
| mNGr-PilZ + Aqs1 YR:SG | mPAO1 with PilZ chromosomally tagged at the N-terminus with mNeonGreen and $His_6$-Aqs1 YR:SG in pHERD30T | This study |
| mNGr-PilZ + Aqs3 | mPAO1 with PilZ chromosomally tagged at the N-terminus with mNeonGreen and $His_6$-Aqs3 in pHERD30T | This study |
| *pilB*::Tn5 + PilB N mut | mPAO1 *pilB* T5 transposon mutant with PilB L154N, L159N, L162N, and I164N in pHERD30T | This study |
| *pilB*::Tn5 + PilB P mut | mPAO1 *pilB* T5 transposon mutant with PilB D156P and V157P in pHERD30T | This study |
| *pilB*::Tn5 + $His_6$-PilB | mPAO1 *pilB* T5 transposon mutant with $His_6$-PilB in pHERD30T | This study |
| *pilB*::Tn5 + $His_6$-PilB D mut | mPAO1 *pilB* T5 transposon mutant with $His_6$-PilB L154D, L159D, L162D in pHERD30T | This study |
| *pilB*::Tn5 + $His_6$-PilB L232R | mPAO1 *pilB* T5 transposon mutant with $His_6$-PilB L232R in pHERD30T | This study |

| | | |
|---|---|---|
| *pilB*::Tn5 + His$_6$-PilB I236R | mPAO1 *pilB* T5 transposon mutant with His$_6$-PilB I236R in pHERD30T | This study |
| *pilB*::Tn5 + His$_6$-PilB D mut L232R | mPAO1 *pilB* T5 transposon mutant with His$_6$-PilB L232R L154D, L159D, L162D in pHERD30T | This study |
| *pilB*::Tn5 + His$_6$-PilB D mut I236R | mPAO1 *pilB* transposon mutant with His$_6$-PilB I236R L154D, L159D, L162D in pHERD30T | This study |
| mPAO1 + Aqs1 YR:SG W45A | mPAO1 with His$_6$-Aqs1 YR:SG and W45A in pHERD30T | This study |
| Δ*gspE* + EV | mPAO1 Δ*gspE* + empty pHERD30T | This study |
| mPAO1 P229A + Aqs1 F19A | mPAO1 with chromosomal P229A mutation with His$_6$-Aqs1 F19A in pHERD30T | This study |
| mPAO1 + pB | mPAO1 with empty pBADGr | This study |
| mPAO1+ pB-Aqs1 YR:SG | mPAO1 with Aqs1 YR:SG in pBADGr | This study |
| mPAO1 + pB-Aqs1 F19A | mPAO1 with Aqs1 F19A in pBADGr | This study |
| ***S. maltophilia* strains** | | |
| WCC C0169 + pB | *S. maltophilia* strain C0169 from the Wright clinical collection with empty pBADGr | This study |
| WCC C0169 + pB-Aqs1 YR:SG | *S. maltophilia* strain C0169 from the Wright clinical collection with Aqs1 YR:SG in pBADGr | This study |
| WCC C0169 + pB-Aqs1 F19A | *S. maltophilia* strain C0169 from the Wright clinical collection with Aqs1 F19A in pBADGr | This study |
| ***A. baylyi* strains** | | |
| ADP1 Parent | ADP1 *comP*$^{T129C}$ | (5) |
| P$_{JExD}$-*aqs1* | ADP1 *comP*$^{T129C}$ Δ*vanAB::kan*$^R$, P$_{JExD}$-*his-aqs1* | This study |
| Δ*pilB* | ADP1 *comP*$^{T129C}$ Δ*pilB* | (5) |
| Δ*tfpB* | ADP1 *comP*$^{T129C}$ Δ*tfpB::gent*$^R$ | This study |
| Δ*pilB* Δ*tfpB* | ADP1 *comP*$^{T129C}$ Δ*pilB* Δ*tfpB::kan*$^R$ | (5) |
| P$_{JExD}$-*aqs1* Δ*pilB* | ADP1 *comP*$^{T129C}$ Δ*pilB* Δ*vanAB::kan*$^R$, P$_{JExD}$-*his-aqs1* | This study |
| P$_{JExD}$-*aqs1* Δ*tfpB* | ADP1 *comP*$^{T129C}$ Δ*tfpB::gent*$^R$ Δ*vanAB::kan*$^R$, P$_{JExD}$-*his-aqs1* | This study |
| **Plasmid list** | | |
| **Plasmid name** | **Characteristics** | **Source** |
| pEX18Gm | Suicide vector for gene replacement | (6) |
| pEX18Gm-Δ*lasR* | *lasR* deletion construct | This study |
| pEX18Gm-Δ*gspE* | *gspE* deletion construct | This study |
| pEX18Gm-PilB F187A | PilB F187A chromosomal knock-in construct | This study |
| pEX18Gm-PilB P228A | PilB P228A chromosomal knock-in construct | This study |
| pEX18Gm-PilB P229A | PilB P229A chromosomal knock-in construct | This study |

| | | |
|---|---|---|
| pEX18Gm-PilA A86C | PilA A86C chromosomal knock-in construct | This study |
| pBADGr | Broad host range arabinose inducible expression vector | (7) |
| pBADGr-Aqs1 YR:SG | Aqs1 Y39S R40G in pBADGr | This study |
| pBADGr-Aqs1 F19A | Aqs1 F19A in pBADGr | This study |
| pHERD30T | Broad host range arabinose inducible expression vector | (8) |
| pHERD30T-Aqs1 | N-terminally $His_6$-tagged Aqs1 in pHERD30T | (2) |
| pHERD30T-Aqs2 | N-terminally $His_6$-tagged Aqs2 in pHERD30T | (2) |
| pHERD30T-Aqs3 | N-terminally $His_6$-tagged Aqs3 in pHERD30T | (2) |
| pHERD30T-Aqs1 YR:SG | N-terminally $His_6$-tagged Aqs1 Y39S R40G in pHERD30T | This study |
| pHERD30T-Aqs1 F19A | N-terminally $His_6$-tagged Aqs1 F19A in pHERD30T | This study |
| pHERD30T-Aqs1 W45A | N-terminally $His_6$-tagged Aqs1 W4A in pHERD30T | This study |
| pHERD30T-Aqs1 YR:SG W45A | N-terminally $His_6$-tagged Aqs1 Y39S, R40G, and W45A in pHERD30T | This study |
| PilB-T18 | mPAO1 full length PilB in pUT18C vector | This study |
| PilB-T25 | mPAO1 full length PilB in pKT25 vector | This study |
| N1D-T18 | mPAO1 PilB N1D in pUT18C vector | This study |
| N1D-T25 | mPAO1 PilB N1D in pKT25 vector | This study |
| N2D/CTD-T18 | mPAO1 PilB N2D and CTD in pUT18C vector | This study |
| N2D/CTD-T25 | mPAO1 PilB N2D and CTD in pKT25 vector | This study |
| LasR-T18 | mPAO1 LasR in pUT18C vector | This study |
| LasR-T25 | mPAO1 LasR in pKT25 vector | This study |
| Aqs1-T18 | mPAO1 Aqs1 in pUT18C vector | (2) |
| Aqs1-T25 | mPAO1 Aqs1 in pKT25 vector | (2) |
| Aqs2-T18 | mPAO1 Aqs2 in pUT18C vector | (2) |
| Aqs2-T25 | mPAO1 Aqs2 in pKT25 vector | (2) |
| Aqs3-T18 | mPAO1 Aqs3 in pUT18C vector | (2) |
| Aqs3-T25 | mPAO1 Aqs3 in pKT25 vector | (2) |
| $PilB^{Gm}$-T18 | *G. metallireducens* PilB in pUT18C vector | This study |
| $PilB^{Gm}$-T25 | *G. metallireducens* PilB in pKT25 vector | This study |
| $PilB^{Ab}$-T18 | *A. baylyi* PilB in pUT18C vector | Howell Lab |
| $PilB^{Ab}$-T25 | *A. baylyi* PilB in pKT25 vector | Howell Lab |

| | | |
|---|---|---|
| TfpB-T18 | *A. baylyi* TfpB in pUT18C vector | Howell Lab |
| TfpB-T25 | *A. baylyi* TfpB in pKT25 vector | Howell Lab |
| MshE-T18 | *V. cholerae* MshE in pUT18C vector | This study |
| MshE-T25 | *V. cholerae* MshE in pKT25 vector | This study |
| CpaF-T18 | *C. crescentus* CpaF in pUT18 vector | Howell Lab |
| CpaF-T25 | *C. crescentus* CpaF in pKNT25 vector | Howell Lab |
| pHERD30T-PilB | Full length wildtype PilB in pHERD30T | (4) |
| pHERD30T-PilB F187A | Full length wildtype PilB F187A in pHERD30T | This study |
| pHERD30T-PilB P228A | Full length wildtype PilB P228A in pHERD30T | This study |
| pHERD30T-PilB P229A | Full length wildtype PilB P229A in pHERD30T | This study |
| pHERD30T-PilB L232A | Full length wildtype PilB L232A in pHERD30T | This study |
| pHERD30T-PilB I236A | Full length wildtype PilB I236A in pHERD30T | This study |
| pETDuet-ΔN1D (ΔD180)-PilB | N-terminally $His_6$-tagged ΔD180-PilB first site and empty second site pETDuet-1 | This study |
| pETDuet-Aqs1 | Empty first site and N-terminally V5-tagged Aqs1 second site pETDuet-1 | This study |
| pETDuet-ΔN1D (ΔD180)-PilB/Aqs1 | N-terminally $His_6$-tagged ΔD180-PilB first site and N-terminally V5-tagged Aqs1 second site pETDuet-1 | This study |
| pETDuet-ΔN1D-PilB F187A/Aqs1 | N-terminally $His_6$-tagged ΔD180-PilB F187A first site and N-terminally V5-tagged Aqs1 second site pETDuet-1 | This study |
| pETDuet-ΔN1D-PilB P228A/Aqs1 | N-terminally $His_6$-tagged ΔD180-PilB P228A first site and N-terminally V5-tagged Aqs1 second site pETDuet-1 | This study |
| pETDuet-ΔN1D-PilB P229A/Aqs1 | N-terminally $His_6$-tagged ΔD180-PilB P229A first site and N-terminally V5-tagged Aqs1 second site pETDuet-1 | This study |
| pETDuet-ΔT148-PilB | N-terminally $His_6$-tagged ΔT148-PilB first site and empty second site pETDuet-1 | This study |
| pETDuet- ΔT148-PilB/Aqs1 | N-terminally $His_6$-tagged ΔT148-PilB first site and N-terminally V5-tagged Aqs1 second site pETDuet-1 | This study |
| pETDuet-$PilB^{Gm}$ | N-terminally $His_6$-tagged ΔN1D-*G. metallireducens* PilB first site and empty second site pETDuet-1 | This study |
| pETDuet-$PilB^{Gm}$/Aqs1 | N-terminally $His_6$-tagged ΔN1D-*G. metallireducens* PilB first site and N-terminally V5-tagged Aqs1 second site pETDuet-1 | This study |
| pETDuet-$PilB^{Ab}$ | N-terminally $His_6$-tagged ΔN1D-*A. baylyi* PilB first site and empty second site pETDuet-1 | This study |
| pETDuet-$PilB^{Ab}$/Aqs1 | N-terminally $His_6$-tagged ΔN1D-*A. baylyi* PilB first site and N-terminally V5-tagged Aqs1 second site pETDuet-1 | This study |
| PilB/PilZ pETDuet | N-terminally $His_6$-tagged PilZ first site and full length PilB second site pETDuet-1 | This study |
| p15TVL-Aqs1 | N-terminally $His_6$-tagged Aqs1 in p15TVL | (2) |
| p15TVL-Aqs1 W45A | N-terminally $His_6$-tagged Aqs1 W45A in p15TVL | This study |
| pCDF-PilB | Full length un-tagged PilB in pCDF | This study |

| | | |
|---|---|---|
| pETDuet-Aqs1/PilB | N-terminally $His_6$-tagged Aqs1 first site and un-tagged full length PilB second site pETDuet-1 | This study |
| pACYDuet-PilZ | N-terminally $His_6$-tagged PilZ first site and empty second site pACYDuet | This study |
| pHERD30T-PilB N mut | PilB L154N, L159N, L162N, and I164N in pHERD30T | This study |
| pHERD30T-PilB P mut | PilB D156P and V157P in pHERD30T | This study |
| pHERD30T-$His_6$-PilB | N-terminally $His_6$-tagged PilB in pHERD30T | This study |
| pHERD30T-PilB D mut | N-terminally $His_6$-PilB L154D, L159D, L162D in pHERD30T | This study |
| pHERD30T $His_6$-PilB L232R | N-terminally $His_6$-tagged PilB L232R in pHERD30T | This study |
| pHERD30T $His_6$-PilB I236R | N-terminally $His_6$-tagged PilB I236R in pHERD30T | This study |
| pHERD30T $His_6$-PilB D mut L232R | N-terminally $His_6$-tagged PilB L154D, L159D, L162D, and L232R in pHERD30T | This study |
| pHERD30T $His_6$-PilB D mut I236R | N-terminally $His_6$-tagged PilB L154D, L159D, L162D, and I236R in pHERD30T | This study |

**Table S2: Primers used in this study.**

| Primers | | |
|---|---|---|
| **Primer name** | **Characteristics** | **Sequence** |
| PaPilB-T18/25_Fwd | PAO1 PilB for cloning into pUT18C and pKT25 | ATATTCTAGAGATGAACGACAGCATCCAACTG |
| PaPilB-T18/25_Rev | PAO1 PilB for cloning into pUT18C and pKT25 | ATATGGTACCTTAATCCTTGGTCACGCGG |
| GmPilB-T18/25_Fwd | *G. metallireducens* PilB for cloning into pUT18C and pKT25 | ATATGGATCCCATGCAGGCCAGCAGACTG |
| GmPilB-T18/25_Rev | *G. metallireducens* PilB for cloning into pUT18C and pKT25 | ATATGGTACCTTAATCGTCGGCCACGGTG |
| Pa PilB-N1D T18/25 | Encodes Pa PilB N1D with ΔD181-C terminus truncation for cloning into pUT18C and pKT25 | ATATGGTACCTTAGTCCGCTTCTGCGCTG |
| Pa PilB-N2D/CTD T18/25 | Encodes Pa PilB N2D and CTD with ΔD180-N terminus truncation for cloning into pUT18C and pKT25 | ATATTCTAGAGGACGCACCTGTAGTACGTTTCG |
| Pa PilB P183A_F | For making PilB P183A mutation | GCGGACGACGCAGCTGTAGTACGTTTCGTC |
| Pa PilB P183A_R | For making PilB P183A mutation | GACGAAACGTACTACAGCTGCGTCGTCCGC |
| Pa PilB F187A_F | For making PilB F187A mutation | CCTGTAGTACGTGCCGTCAACAAGATGCTG |
| Pa PilB F187A_R | For making PilB F187A mutation | CAGCATCTTGTTGACGGCACGTACTACAGG |
| Pa PilB P183A_F | For making PilB P228A mutation | CTCCACGAAGTGGCCAAGGCGCCGATCCAGTTGGCC |

| | | |
|---|---|---|
| Pa PilB P183A_R | For making PilB P228A mutation | GGCCAACTGGATCGGCGCCTTGGCCACTTCGTGGAG |
| Pa PilB F187A_F | For making PilB P229A mutation | CACGAAGTGGCCAAGCCGGCGATCCAGTTGGCC |
| Pa PilB F187A_R | For making PilB P229A mutation | GGCCAACTGGATCGCCGGCTTGGCCACTTCGTG |
| Pa PilB L232A_F | For making PilB L232A mutation | GTCGCCAAGCCGCCGATCCAGGCCGCCAGTCGTATCTCTGCTCGTC |
| Pa PilB L232A_R | For making PilB L232A mutation | GACGAGCAGAGATACGACTGGCGGCCTGGATCGGCGGCTTGGCGAC |
| Pa PilB I236A_F | For making PilB L236A mutation | GTCGCCAAGCCGCCGATCCAGTTGGCCAGTCGTGCCTCTGCTCGTCTCAAGGTA |
| Pa PilB I236A_R | For making PilB L236A mutation | TACCTTGAGACGAGCAGAGGCACGACTGGCCAACTGGATCGGCGGCTTGGCGAC |
| PilBPa F187A chrom mut_F | For knocking F187A onto the chromosome | GCACCTGTAGTACGCGCCGTCAACAAGATGCTGCT |
| PilBPa F187A chrom mut_R | For knocking F187A onto the chromosome | AGCAGCATCTTGTTGACGGCGCGTACTACAGGTGC |
| PilB F187A Ups Fwd | For amplifying region upstream of PilB F187 | ATATTCTAGACTACTCGACGAAAAGACCGCC |
| PilB F187A Dns Rev | For amplifying region downstream of PilB F187 | ATATGGTACCGTTCAGGCCGGTGTATAGCG |
| PilBPa P228A chrom mut_F | For knocking P228A onto the chromosome | GTCGCCAAGGCGCCGATCCAGTTGGCCAGT |
| PilBPa P228A chrom mut_R | For knocking P228A onto the chromosome | ACTGGCCAACTGGATCGGCGCCTTGGCGAC |
| PilB P228A Ups Fwd | For amplifying region upstream of PilB P228 | ATATTCTAGAGAGCAGTTCGGCATCGCCTATT |
| PilB P228A Dns Rev | For amplifying region downstream of PilB P228 | ATATGGTACCCGCCGACCATGATCACATCC |
| PilBPa P229A chrom mut_F | For knocking P229A onto the chromosome | CACGAAGTCGCCAAGCCGGCGATCCAGTTGGCCAGTC |
| PilBPa P229A chrom mut_R | For knocking P229A onto the chromosome | GACTGGCCAACTGGATCGCCGGCTTGGCGACTTCGTG |
| PilB P229A Ups Fwd | For amplifying region upstream of PilB P229 | ATATTCTAGACCGAGCAGTTCGGCATCG |
| PilB P229A Dns Rev | For amplifying region downstream of PilB P229 | ATATGGTACCTCTCGCCGACCATGATCACA |
| PilB N2D/CTD_Fwd | Encodes Pa PilB N2D and CTD with ΔD180-N terminus truncation for cloning into pET28b | ATATGCTAGCGACGCACCTGTAGTACGTTTCGTC |
| Pa PilB_Rev | For cloning PilB into pET28b | ATATGAATTCTTAATCCTTGGTCACGCGGTT |
| Aqs1 pBAD_Fwd | For cloning Aqs1 into pBADGr | ATATGAATTCATGACAAACACCGACCTCAAAC |
| Aqs1 pBAD_Rev | For cloning Aqs1 into pBADGr | ATATAAGCTTTCATTCATGGTCGCCCC |
| LasR-T18/25_Fwd | For cloning full length PAO1 LasR into pUT18C and pKT25 | ATATGGATCCCATGGCCTTGGTTGACG |

| | | |
|---|---|---|
| LasR-T18/25_Rev | For cloning full length PAO1 LasR into pUT18C and pKT25 | ATATGGTACCTCAGAGAGTAATAAGACCCAAATTAAC |
| PilB N2DDuet_F | Encodes Pa PilB N2D and CTD with ΔD180-N terminus truncation for cloning into pETDuet-1 first site | ATATGAATTCGGACGCACCTGTAGTACGTTTCGTC |
| PilB N2DDuet_R | Encodes Pa PilB N2D and CTD with ΔD180-N terminus truncation for cloning into pETDuet-1 first site | ATATGCGGCCGCTTAATCCTTGGTCACGCGGTT |
| Aqs1 V5 Tag pETDuet_F | For cloning Aqs1 with a V5 tag into pETDuet-1 | ATATCAATTGGGGCAAACCGATTCCGAACCCGCTGCTGGGCCTGGATAGCACCATGACAAACACCGACCTCAAAC |
| Aqs1 V5 Tag pETDuet_R | For cloning Aqs1 with a V5 tag into pETDuet-1 | ATATGGTACCTCATTCATGGTCGCCCC |
| PilB pHERD_Fwd | For cloning PilB into pHERD30T | ATATGAGCTCGCGATTCCTTCCCCATGA |
| PilB pHERD_Rev | For cloning wild-type PilB into pHERD30T | ATATTCTAGATTAATCCTTGGTCACGCGG |
| PilA A86C Fwd | For mutating PAO1 PilA A86C | GGCGTCGAGCCGGATTGTAACAAGTTGGGTGTA |
| PilA A86C Rev | For mutating PAO1 PilA A86C | TACACCCAACTTGTTACAATCCGGCTCGACGCC |
| PilA A86 Ups_Fwd | PilA A86 primer for upstream region amplification | ATATGAATTCCGCTCAGTTGGATGCTGTC |
| PilA A86 Dns_Rev | PilA A86 primer for downstream region amplification | ATATTCTAGAGCCAAGCTGGAAGCTTCC |
| LasR UpS_F | For making a *lasR* deletion upstream region | ATATGAATTCGGGGACCAGGTGTGACTG |
| LasR UpS_R | For making a *lasR* deletion upstream region | ATATGGATCCAGGATGGCGCTCCACTC |
| LasR DnS_F | For making a *lasR* deletion downstream region | ATATGGATCCCGGAAGTTCGGTGTGACC |
| LasR DnS_R | For making a *lasR* deletion downstream region | ATATAAGCTTCCATCGATTTCCATCTCGTC |
| GspE Ups_F | For making a *gspE* deletion upstream region | ATATGAATTCCTGCGGATCGACTCGTGG |
| GspE Ups_R | For making a *gspE* deletion upstream region | ATATTCTAGAGCGAAGCTGAACGGCAGAC |
| GspE Dns_F | For making a *gspE* deletion downstream region | ATATTCTAGACAAGGTGCTGGAAGGCGTC |
| GspE Dns_R | For making a *gspE* deletion downstream region | ATATAAGCTTTGCTCGGTGTAGTCGGCAAG |
| mNGr-PilZ_F1 | For cloning a PilZ mNeonGreen N-terminal fusion | ATATGAATTCGGCAGGAACCTGCATGGTGAGCAAGGGCGAGGAG |
| mNGr-PilZ_R1 | For making an mNeonGreen-PilZ N-terminal fusion construct - encodes 5G linker | ACCACCACCACCACCCTTGTACAGCTCGTCCATGCC |
| mNGr-PilZ_F2 | For cloning a PilZ mNeonGreen N-terminal fusion | GGTGGTGGTGGTGGTATGAGTTTGCCACCCAATCTG |
| mNGr-PilZ_R2 | For cloning a PilZ mNeonGreen N-terminal fusion | ATATAAGCTTTTACATCGTGTGGGTCGGC |

| | | |
|---|---|---|
| PilB F187ADuet_F | For cloning PilB F187A into pETDuet-1 | ATATGAATTCGGACGCACCTGTAGTACGTGCCGTC |
| PilB linker N mut_F | Mutates all L and I residues between A147-179 to N | AACGCAGGCAACGACGATGTCGACAACGAGGGAAATGATAAC |
| PilB linker N mut_R | Mutates all L and I residues between A147-179 to N | GTTATCATTTCCCTCGTTGTCGACATCGTCGTTGCCTGCGTT |
| PilB linker P mut_F | Mutates PilB D156 and V157 to P | CGCAGGCCTCGACCCTCCCGACCTCGAGGGACTGG |
| PilB linker P mut_R | Mutates PIlB D156 and V157 to P | CCAGTCCCTCGAGGTCGGGAGGGTCGAGGCCTGCG |
| MshE-T18/25_F | For cloning *mshe* into BACTH T18/T25 vectors | ATATTCTAGAGGTGCCAATTAATAAACTGCGTAAAC |
| MshE-T18/25_R | For cloning *mshe* into BACTH T18/T25 vectors | ATATGAATTCCTACAGATAAATCGGTTCAACCAA |
| N-His PilB 30T_F | For cloning PilB with an N-terminal His-tag into pHERD30T | ATATGAGCTCATTCCTTCCCCATGCATCATCATCATCATCACAACGACAGCATCCAA |
| PilB I236R_F | For making a PilB I236R mutation | ATCCAGTTGGCCAGTCGTCGCTCTGCTCGTCTCAAGG |
| PilB I236R_R | For making a PilB I236R mutation | CCTTGAGACGAGCAGAGCGACGACTGGCCAACTGGAT |
| PilB L232R_F | For making a PilB L232R mutation | GCCAAGCCGCCGATCCAGCGGGCCAGTCGTATCTCTG |
| PilB L232R_R | For making a PilB L232R mutation | CAGAGATACGACTGGCCCGCTGGATCGGCGGCTTGGC |
| PilB L154D_F | For making a PilB L154D mutation | CCGACGGTCTCGCAGGCGACGACGATGTCGACCTCG |
| PilB L154D_R | For making a PilB L154D mutation | CGAGGTCGACATCGTCGTCGCCTGCGAGACCGTCGG |
| PilB L159D_F | For making a PilB L159D mutation | CTCGACGATGTCGACGACGAGGGACTGGATATC |
| PilB L159D_R | For making a PilB L159D mutation | GATATCCAGTCCCTCGTCGTCGACATCGTCGAG |
| PilB L162D_F | For making a PilB L162D mutation | GTCGACCTCGAGGGAGACGATATCGGTAGCGCGGAC |
| PilB L162D_R | For making a PilB L162D mutation | GTCCGCGCTACCGATATCGTCTCCCTCGAGGTCGAC |
| PilB Linker D mut_F | For making a PilB L154D, L159D, and L162D mutant | CTCGCAGGCGACGACGATGTCGACGACGAGGGAGACGATATCGGTAGCGCGGAC |
| PilB Linker D mut_F | For making a PIlB L154D, L159D, and L162D mutant | GTCCGCGCTACCGATATCGTCTCCCTCGTCGTCGACATCGTCGTCGCCTGCGAG |

| | | |
|---|---|---|
| PilB linker T18/25_F | For cloning PilB linker T148-D180 into T18 and T25 | ATATTCTAGAGACCGACGGTCTCGCAGGC |
| PilB linker T18/25_R | For cloning PilB linker T148-D180 into T18 and T25 | ATATGAATTCTTAGTCCGCTTCTGCGCTGG |
| PilB Ab N2D/CTD Duet_F | For cloning *A. baylyi* PilB from K188 into pETDuet-1 | ATATGAATTCGAAACTTAAAGACGAAGCACCTATTG |
| PilB Ab N2D/CTD Duet_R | For cloning *A. baylyi* PilB from K188 into pETDuet-1 | ATATAAGCTTTTATTCACTGGTTACACGATTAATTTC |
| PilB Gm N2D/CTD Duet_F | For cloning *G. metallireducens* PilB from D180 into pETDuet-1 | ATATGAATTCGGACGCTCCGGTCGTGAAG |
| PilB Gm N2D/CTD Duet_R | For cloning *G. metallireducens* PilB from D180 into pETDuet-1 | ATATCTGCAGTTAATCGTCGGCCACGGT |
